# Sulfoquinovose is exclusively metabolized by the gut microbiota and degraded differently in mice and humans

**DOI:** 10.1101/2025.01.22.634256

**Authors:** Julia Krasenbrink, Buck T. Hanson, Anna S. Weiss, Sabrina Borusak, Tomohisa Sebastian Tanabe, Michaela Lang, Georg Aichinger, Bela Hausmann, David Berry, Andreas Richter, Doris Marko, Marc Mussmann, David Schleheck, Bärbel Stecher, Alexander Loy

## Abstract

**Background:** Sulfoquinovose (SQ) is a green-diet-derived sulfonated glucose and a selective substrate for few human gut bacteria. Complete anaerobic SQ degradation via interspecies metabolite transfer to sulfonate-respiring bacteria produces hydrogen sulfide, which has dose- and context-dependent health effects. Here, we studied potential SQ degradation by the mammalian host and the impact of SQ supplementation on human and murine gut microbiota diversity and metabolism.

**Results:** ^13^CO_2_ breath tests with germ-free C57BL/6 mice gavaged with ^13^C-SQ were negative. Also, SQ was not degraded by human intestinal cells *in vitro*, indicating that SQ is not directly metabolized by mice and humans. Addition of increasing SQ concentrations to human fecal microcosms revealed dose-dependent responses of the microbiota and corroborated the relevance of *Agathobacter rectalis* and *Bilophila wadsworthia* in cooperative degradation of SQ to hydrogen sulfide via interspecies transfer of 2,3-dihydroxy-1-propanesulfonate (DHPS). Similar to the human gut microbiome, the genetic capacity for SQ or DHPS degradation is sparsely distributed among bacterial species in the mouse gut. *Escherichia coli* and *Enterocloster clostridioformis* were identified as primary SQ degraders in the mouse gut. SQ and DHPS supplementation experiments with conventional laboratory mice and their intestinal contents showed that SQ was incompletely catabolized to DHPS. Although some *E. clostridioformis* genomes encode an extended sulfoglycolytic pathway for both SQ and DHPS fermentation, SQ was only degraded to DHPS by a mouse-derived *E. clostridioformis* strain.

**Conclusions:** Our findings suggest that SQ is solely a nutrient for the gut microbiota and not for mice and humans, emphasizing its potential as a prebiotic. SQ degradation by the microbiota of conventional laboratory mice differs from the human gut microbiota by absence of DHPS degradation activity. Hence, the microbiota of conventional laboratory mice does not fully represent the SQ metabolism in humans, indicating the need for alternative model systems to assess the impact of SQ on human health. This study advances our understanding of how individual dietary compounds shape the microbial community structure and metabolism in the gut and thereby potentially influence host health.

## Introduction

The sulfonated glucose sulfoquinovose (6-deoxy-6-sulfo-D-glucose, SQ) is an abundant sulfur molecule in the global sulfur cycle. As the polar headgroup of the sulfonolipid sulfoquinovosyldiacylglycerol (SQDG), an estimated 10 billion tons of SQ is synthesized annually by photosynthetic organisms such as plants, algae, and cyanobacteria [1]. SQDG is located in thylakoid membranes and maintains the activity and structural integrity of photosystem II [2–5]. Sulfoglycolytic microorganisms can cleave SQ from SQDG or other SQ glycosides via a sulfoquinovosidase. Two types of sulfoquinovosidase enzymes are known.

The YihQ sulfoquinovosidase belongs to glycoside hydrolase family 31 [6] and the NAD^+^-dependent SqgA sulfoquinovosidase belongs to glycoside hydrolase family 188 [7]. To date, six microbial SQ degradation pathways have been described. In two pathways, SQ is catabolized aerobically to sulfite by individual organisms using an alkanesulfonate monooxygenase [8] or an iron- and alpha-ketoglutarate-dependent alkanesulfonate dioxygenase [9]. The other pathways require syntrophic interspecies metabolite transfer of C_2_- or C_3_-organosulfonates for complete degradation. The sulfo-Embden-Meyerhof-Parnas pathway generates the C_3_-organosulfonates 2,3-dihydroxypropane-1-sulfonate (DHPS) or 3-sulfolactate (SL) during aerobic or anaerobic growth [10,11]. The sulfo-Entner-Doudoroff-pathway degrades SQ aerobically to SL [12]. The sulfofructose-transaldolase pathway produces SL aerobically and SL or DHPS anaerobically [13,14]. The sulfo-transketolase pathway degrades SQ to two molecules of the C_2_-organosulfonates isethionate or, putatively, sulfoacetate [11]. In these pathways, the residual carbon skeleton (either C_3_ or C_2_) enters the central carbon metabolism while organosulfonates are excreted.

DHPS is a key metabolite of SQ-degrading human gut bacteria (Figure 1 a) [15]. Four pathways for anaerobic DHPS degradation are currently known (Figure 1 b). Two of these pathways are utilized by select sulfite-reducing bacteria (*Desulfovibrio*, *Bilophila*) and result in the intracellular formation of sulfite, which is subsequently respired to hydrogen sulfide (H_2_S). In *Desulfovibrio* strain DF1, DHPS is degraded to pyruvate and sulfite via 3-sulfolactaldehyde and SL [16]. In *Bilophila wadsworthia*, DHPS is cleaved to hydroxyacetone and sulfite [15,17]. Some fermenting bacteria, such as *Klebsiella oxytoca* and *Hungatella hathewayi*, can degrade DHPS by reducing it to 3-hydroxypropane-1-sulfonate (3-HPS) via sulfopropionaldehyde in a third pathway [17]. In a fourth, bifurcated pathway in *Enterococcus gilvus*, DHPS is converted to both 3-HPS and 3-sulfopropionate [18]. In the human gut, *B. wadsworthia* may use 3-sulfopropionate, but not 3-HPS, as a terminal electron acceptor resulting again in H_2_S formation [18]. The strictly anaerobic mouse gut bacterium *Taurinivorans muris* encodes another potential DHPS degradation pathway, but did not degrade DHPS in growth tests [19]. However, this pathway was thus far only demonstrated in aerobic bacteria, which have the sulfopropanediol-3-dehydrogenase HpsN to oxidize (*R*)-DHPS to (*R*)-sulfolactate [20], which is then cleaved to pyruvate and sulfite by sulfolactate-sulfolyase SuyAB [21].

**Figure 1:**
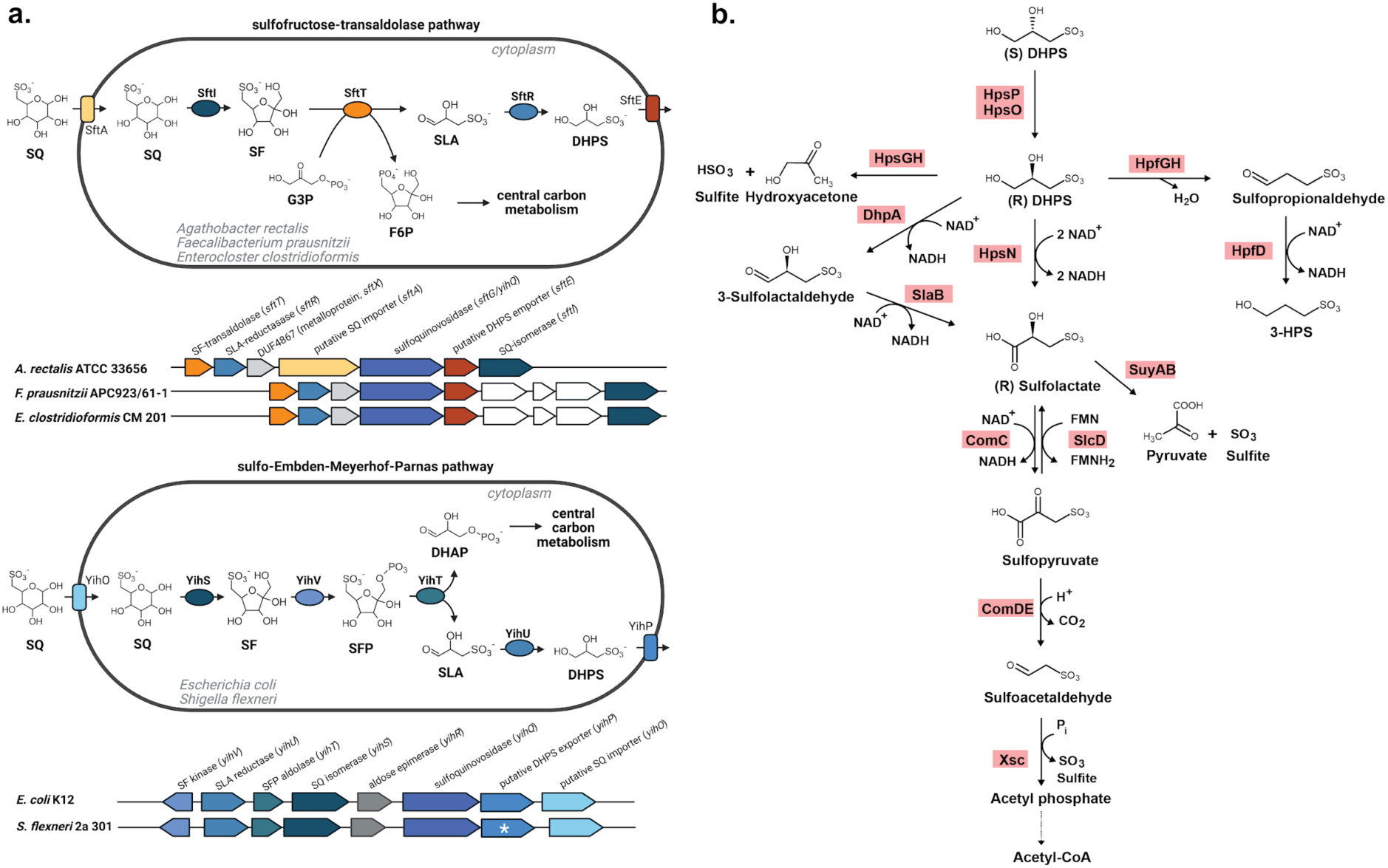
Prevalent SQ degradation pathways in the human gut and overview of enzymatic DHPS degradation reactions. SQ is degraded to DHPS via the sulfofructose-transaldolase pathway in *Agathobacter rectalis* (GCF_000020605.1), *Faecalibacterium prausnitzii* (GCF_003287495.1), and *Enterocloster clostridioformis* (GCF_000371405.1) or via the sulfo-Embden-Meyerhof-Parnas pathway in *Escherichia coli* (GCF_000005845.2) and *Shigella flexneri* (GCF_000006925.2) (created with biorender.com) (a). Genes depicted in white have no predicted function. The white asterisk indicates that the *yihP* homolog of *S. flexneri* contains a frameshift and is thus a potential pseudogene. Overview of known DHPS degradation pathways (b). HpsP, DHPS-2-dehydrogenase; HpsO, DHPS-2-dehydrogenase; HpsG, DHPS-sulfolyase; HpsH, DHPS-sulfolyase activating enzyme; HpfG, DHPS-dehydratase; HpfH, DHPS-dehydratase activating enzyme; HpfD, sulfopropionaldehyde reductase; HpsN, DHPS-1-dehydrogenase; DhpA, dehydrogenase; SlaB, dehydrogenase; SuyAB sulfolactate-sulfolyase; ComC, sulfolactate dehydrogenase; SlcD, sulfolactate dehydrogenase; ComDE, 3-sulfopyruvate decarboxylase; Xsc, sulfoacetaldehyde acetyltransferase. SQ, sulfoquinovose; SF, 6-deoxy-6-sulfofructose; SLA, 3-sulfolactaldehyde; DHPS, 2,3-dihydoxypropane-1-sulfonate; G3P, glycerinaldehyde-3-phosphate; F6P, fructose-6-phosphate; SFP, 6-deoxy-6-sulfofructose phosphate; DHAP, dihydroxyacetone phosphate.

As a component of edible green plants and algae, SQDG is part of the human and animal diet. While the capability of mammalian cells to metabolize SQ remains unclear, SQ is a selective substrate for a few but abundant human gut bacteria [15]. *Agathobacter rectalis* (formerly *Eubacterium rectale*) and *B. wadsworthia* were identified as main primary and secondary SQ degraders in human fecal microcosms, respectively. Nearly half of the high-quality *A. rectalis* genomes from a human microbiome database [22] encoded SQ degradation via the sulfofructose-transaldolase pathway, though this potential was rare among other human gut bacteria, with only 0.5% of species-level genome bins containing the pathway. Despite its limited genomic prevalence, the sulfofructose-transaldolase pathway ranked among the top third most abundant microbial pathways in fecal metatranscriptome datasets of four human cohorts. This highlights SQ degradation as a core function of the human gut microbiome, with *A. rectalis* and *Faecalibacterium prausnitzii* contributing most to its transcription [15]. Primary SQ degraders are generally associated with beneficial host effects [23,24] and thus SQ is a candidate prebiotic. However, excessive stimulation of secondary, H_2_S-producing SQ degraders may be harmful as H_2_S has dose-dependent impacts on host health [25]. At low nano-to micromolar concentrations, H_2_S has predominantly beneficial effects [26]. H_2_S serves as an energy source for mitochondria, as a cellular antioxidant, and as a gasotransmitter in mammals [27]. Additionally, H_2_S may prevent colonization by pathogens by inhibiting their aerobic respiration [28]. At high micro-to millimolar concentrations produced by the gut microbiota [26], however, H_2_S may disrupt disulfide bonds in the intestinal mucosal layer, thus enabling bacterial penetration [29], and can lead to cell death by inhibiting cytochrome *c* [30]. Overabundance of H_2_S-producing bacteria, such as *B. wadsworthia*, is considered as a potential driver of gut inflammation [29,31,32].

Here, we assessed the prebiotic potential of SQ by studying its degradation and metabolism in humans and mice using both human and mouse gut content microcosm experiments, as well as *in vivo* interventions in conventional and germ-free mice. We elucidated the role of mammalian cells in SQ metabolism through *in vitro* cultures and stable isotope-tracing in germ-free mice. We show that SQ is not degraded by the host, establish a minimum SQ concentration that impacts the human gut microbiota, and uncover fundamental differences in the bacterial SQ metabolism between wild type mice and humans.

## Materials and Methods

### Human cell culture

HT-29, a human colorectal adenocarcinoma cell line (German Collection of Microorganisms and Cell Cultures, Germany), was cultured in Dulbecco’s Modified Eagles Medium (DMEM), which contains 25 mM glucose, supplemented with 10% (v/v) fetal calf serum and 1% (v/v) penicillin/streptomycin at 37°C and 5% CO_2_ under humidified conditions. All components were purchased from Invitrogen^TM^ Life Technologies (Germany). Cells were regularly monitored for mycoplasma contamination.

To investigate whether human intestinal cells degrade SQ, HT-29 cells were grown in DMEM or in glucose-free DMEM either supplemented with 25 mM glucose, 25 mM SQ, or left unamended. Supernatants were sampled after 0, 1, 6, 24, and 48 h and stored at −20°C for metabolite analysis by capillary electrophoresis.

To study the potential inhibitory effect of SQ on the growth of HT-29 cells, 7,500 cells were seeded per well and incubated in DMEM medium supplemented with glucose or SQ at three concentrations (2.5 mM, 25 mM, 250 mM). In three independent experiments, duplicate incubations (n = 6) were performed for all treatments. Protein production was measured as a proxy for growth after 3, 5, and 7 days of incubation using sulforhodamine B colorimetric assays as described previously [33]. Briefly, plates were washed with 1x PBS twice and cells were fixed with a 5% trichloroacetic acid solution. Plates were dried overnight at room temperature. Then, the plates were stained with 0.4% (w/v) sulforhodamine B for 1 h. After washing twice with water and twice with 1% (v/v) acetic acid, the stain was eluted using a 10 mM Tris buffer (pH 10). Absorbance was measured at 570 nm and normalized to the solvent control (1% v/v DMSO) to assess the impact of treatments on cell growth.

### Germ-free mouse experiment and ^13^CO_2_ breath test

Six germ-free male C57BL/6 mice (6–20 weeks old), were housed under sterile conditions in flexible film isolators (North Kent Plastic Cages, England) or in Han-gnotocages (ZOONLAB GmbH, Germany). Mice were kept on a twelve-hour light-dark cycle at 20.5-23.5°C and 45-65% humidity, with autoclaved water and Mouse-Breeding complete feed (Ssniff, Germany) provided *ad libitum*. Mice were scored twice daily for their health status, and sterility was checked monthly during housing, before the experiment, and during the experiment using qPCR with universal bacterial 16S rRNA gene primers [34,35]. Mice were randomly assigned to two treatment groups and kept in separate cages. The administered substrates were either 10 mg of fully ^13^C-labeled D-glucose (Sigma-Aldrich, Germany) or 10 mg of fully ^13^C-labeled SQ, which was synthesized from fully ^13^C-labeled D-glucose following an established protocol [36]. Each treatment group received their substrate dissolved in 100 µl sterile water (Ampuwa; Fresenius Kabi, Germany) via gavage. Respired air from individual mice was sampled for CO_2_ stable isotope analysis prior to gavage (baseline) and at 1, 3, 6, 9, 12, and 24 h post-gavage. For breath air sampling, individual mice were placed in an airtight one-liter container sealed with a lid fitted with a rubber stopper for approximately 2 minutes. Separate containers were used for each treatment group to avoid cross-contamination between groups. Gas samples (15 ml) were taken at 30 and 90 seconds and injected entirely into pre-vacuumed exetainers (12 ml) creating an overpressure to prevent leakage from ambient air into the exetainers. Ambient air samples were taken before the experiment for blank measurements. If feasible, fecal samples were collected to validate sterility of mice and for metabolite analysis. ^13^CO_2_ concentrations were measured in the gas samples using isotope ratio mass spectrometry (GasBench II headspace sampler connected to a Delta V Advantage isotope mass spectrometer; Thermo Fisher Scientific, Germany). ^13^CO_2_ production rates per minute were calculated from the differences in ^13^CO_2_ (atom percent excess) between the two samples taken at 30 s and 90 s.

### Anoxic *in vitro* gut microcosm incubations and pure culture growth tests

All anoxic incubations and growth tests were performed in an anaerobic tent (Coy Laboratory Products, United States) operated with a gas mix of 85% N_2_, 10% CO_2_ and 5% H_2_. The H_2_ levels in the tent atmosphere ranged between 1.5 and 2.5%.

Fecal samples were collected from five human individuals, who had not taken antibiotics in the three months before sampling. Samples were stored at 4°C for maximally 24 h before processing. All samples were pooled, and 3.9 g of fecal mass was homogenized in oxygen-free 30 mM bicarbonate buffer (pH 6.8). This fecal slurry (1.32 mg/ml) was distributed into 20 ml Hungate tubes (final volume: 15 ml) and sealed with butyl-rubber stoppers and screw-on caps. Triplicate microcosms were amended with 10 mM, 1 mM, 0.1 mM, or 0.01 mM SQ (MCAT GmbH, Germany). One microcosm was left unamended (negative control). Tubes were incubated at 37°C and subsamples (1 ml) were collected after 0, 3, 6, 9, 12, 16, and 24 h. From each subsample, 20 µl was used for sulfide quantification. The remaining subsample was centrifuged at 11,000 x g for 5 min at 4°C. Biomass-free supernatants were stored at −20°C for subsequent capillary electrophoresis. Cell pellets were frozen at −80°C for DNA extraction.

In two independent experiments, we sampled intestinal contents of conventional C57BL/6 wild type mice to create mouse gut content slurries. Mice were housed under specific-pathogen-free conditions with standard chow diet (Ssniff V1534, Ssniff, Germany) and water *ad libitum*, in compliance with Federation of European Laboratory Animal Science Associations guidelines [37]. Specific pathogen-free status was confirmed through regular screening for viral, bacterial, mycoplasmal, and parasitic agents such as murine norovirus, mouse hepatitis virus, *Helicobacter* species, *Salmonella* species, *Streptococcus* species, protozoa, and helminths. After euthanasia, mice were dissected in an anaerobic chamber.

In the first experiment, ten mice (4 females, 6 males, 8–10 weeks old) were dissected, and their small intestinal, cecal and large intestinal contents were pooled and homogenized with oxygen-free 1x PBS (pH 7.4). The gut content slurry (5.25 mg/ml) was distributed into 20 ml Hungate tubes (final volume: 10 ml). Mouse gut microcosms were left unamended or amended with 10 mM SQ or 10 mM DHPS. All treatments were performed in triplicate. Hungate tubes were sealed with butyl-rubber stoppers and screw-on caps and incubated at 37°C. Subsamples were collected at 0, 24, 48, and 96 h, and sulfide was quantified in 20 µl of subsample.

In the second experiment, eight mice (4 females, 4 males, 8–10 weeks old) were dissected, and their small intestinal (2.2 g) and cecal contents (5.9 g) were separately pooled. Small intestine and cecal contents were homogenized with oxygen-free 1x PBS (pH 7.4) and transferred into 20 ml Hungate tubes (final volume: 15 ml) with a concentration of 4.9 mg/ml and 5.15 mg/ml of small intestinal and cecal content, respectively. Both sets of microcosms were left unamended or amended with 10 mM SQ, and sealed with butyl-rubber stoppers and screw-on lids for incubation at 37°C. All treatments were performed in triplicate. Subsamples (1 ml) were taken at 0, 24, 120, and 220 h. In both experiments, subsamples were centrifuged at 11,000 x g for 5 min at 4°C. Pellets were stored for DNA extraction at −80°C and biomass-free supernatants were stored at −20°C for metabolite analyses.

*Enterocloster clostridioformis* YL32 (DSM 26114) and *Faecalibaculum rodentium* ALO17 (DSM 103405) were each grown in triplicate cultures in 100% and 30% brain heart infusion broth with or without 10 mM SQ at 37°C in gas-tight Hungate tubes. Anoxic and autoclaved ultrapure water was used for dilution of the medium. Growth was monitored by measuring optical density at 600 nm. Identities of grown strains were confirmed by direct Sanger sequencing (Microsynth GmbH, Austria) of 16S rRNA gene amplicons.

### Conventional wild type mouse experiment

Fifteen conventional C57BL/6 wild type mice (6-8 weeks) were bred and housed under specific pathogen-free conditions in accordance with the guidelines of the Federation of European Laboratory Animal Science Associations [37]. As aforementioned, specific pathogen-free status was confirmed through regular screening. Mice were maintained with *ad libitum* water and autoclaved mouse chow (Ssniff V1534, Ssniff, Germany). The day/night cycle was 14/10 hours, and environmental conditions were maintained at 20–24°C with 45–65% humidity. After one week of acclimatization, mice were randomly assigned to three groups (two males and three females each) with no significant differences in weight between groups. Two treatment groups were gavaged with 1 mg or 10 mg SQ in 100 µl aqueous solutions, while the third, control group was gavaged with 100 µl autoclaved tap water. Fecal samples were collected 0, 3, 6, 9, 12, and 24 h post-gavage. General discomfort of treated mice was assessed by observing them after gavage and throughout the experiment for signs of reduced mobility, scruffy fur, hunched posture or changes in facial expression. Mice were sacrificed and dissected after 24 h. Fecal samples and intestinal contents were immediately stored at −80°C for metabolite analysis and 16S rRNA gene amplicon sequencing.

### Metabolite analysis

For sulfide quantification, 20 µl subsamples were added to 300 µl 2 g/l zinc acetate solution in the anaerobic tent. Fixed samples were quantified colorimetrically using the Infinite M Nano+ microplate reader (Tecan, Austria), as described previously [38].

Capillary electrophoresis was used to quantify metabolites in the supernatants of human cell cultures, in gut microcosms, in bacterial pure cultures, and in the fecal samples of germ-free and conventional mice. Concentrations of SQ, SL, DHPS, lactate, and the short-chain fatty acids (SCFAs) formate, succinate, acetate, propionate, butyrate, and valerate were quantified using a 7100 capillary electrophoresis system (Agilent, United States). Mouse fecal samples were prepared by adding 40 µl of ultrapure water to 0.01 g of feces. After mixing, samples were centrifuged at 11,000 x g for 10 min at 4°C and supernatants were stored at −20°C. For analysis, culture supernatants or metabolite extracts from fecal samples were diluted 1:5-1:20 in ultrapure water containing 0.1 mM of an internal standard. Caproate, valerate or succinate was used as an internal standard and was selected based on the sample background. Analyte standard mixtures were prepared in concentrations of 0.025 mM, 0.05 mM, 0.1 mM, 0.25 mM, 0.5 mM and 1 mM. An organic acid buffer (Agilent, United States) was used as a running buffer. The instrument’s bare fused silica capillary (75 µm inner diameter, 72 cm effective length; Agilent, United States) was flushed within the system for four minutes as part of the preconditioning process before each measurement. Samples were then injected at 50 mbar for five seconds. Measurements were performed at –25 kV, 100 µA, and 6 W.

SQ and its degradation products DHPS, SL, isethionate, 3-HPS, 3-sulfopropionate, and sulfoacetate in mouse gut content microcosms, *E. clostridioformis* pure cultures, and in fecal samples of germfree and conventional mice were additionally quantified using liquid chromatography-mass spectrometry. Samples were prepared by centrifuging at 18,000 x g for 5 minutes at 4°C. The supernatants were then diluted 1:10 with ultrapure water and mixed at a 7:3 (v/v) ratio with ≥ 99.9% acetonitrile containing 6.7 mM para-toluenesulfonate as internal standard. Samples were quantified using a LCMS-2020 single quadrupole mass spectrometer system (Shimadzu, Japan) consisting of LC-40B XR pumps, a SIL-40C XR autosampler, a CTO-40S column oven (heated to 30°C) and a MS-2020 electrospray ionization mass spectrometer. Organosulfonate separation was carried out on an iHILIC-Fusion(+) column (100 × 2.1 mm, 3.5 µm, 100 Å), as described previously [39].

### 16S rRNA gene amplicon sequencing and real-time PCR

DNA extraction and 16S rRNA gene amplicon sequencing was performed by the Joint Microbiome Facility of the Medical University of Vienna and the University of Vienna under the project numbers JMF-2012-06 (human fecal microcosms), JMF-2108-01 (conventional mouse experiment), JMF-2212-09 (mouse gut content microcosms). DNA was extracted with the QIAamp Fast DNA Stool Mini Kit according to the manufacturer’s protocol (Qiagen GmbH, Austria). Extraction contamination was controlled for using blank extractions containing kit-provided buffers alone. An established two-step PCR-barcoding method (H_515F_mod-5’-(head) GCT ATG CGC GAG CTG C - GTG YCA GCM GCC GCG GTA A and H_806R_mod-5’-(head) TAG CGC ACA CCT GGT A - GGA CTA CNV GGG TWT CTA AT) was used for 16S rRNA gene amplicon sequencing of bacteria and archaea [40]. Prior to preparing the sequencing libraries, barcoded amplicon libraries were normalized with the SequalPrep normalization plate kit and subsequently pooled (TrueSeq Nano Kit; Illumina, United States) [40,41]. Amplicon sequencing was performed on the MiSeq (V3, 600 cycles; Illumina, United States) and results were analyzed according to the procedures outlined previously [40]. Amplicon sequence variants (ASVs) were inferred using the DADA2 workflow [42] and taxonomically classified with the SILVA database (release 138). Sequencing libraries were rarified and analyzed with the software packages phyloseq (v1.42.0) [43] and vegan (v2.6-4) [44] for RStudio (rstudio.com, R version 4.2.2). Principal coordinate analysis was performed using a Bray-Curtis dissimilarity matrix computed with phyloseq. DESeq2 (v1.38.3) was used to perform differential abundance analyses between treatment groups and time points [45].

In the human fecal microcosm experiment, ASV abundance at each time point and treatment was compared to two controls: Its corresponding baseline at 0 h, and to the abundance in the 0.01 mM SQ incubation at the same time point. Notably, the 0.01 mM SQ incubation was used as control because (i) metabolite data showed no significant differences compared to the unamended control and (ii) only one control incubation was sequenced. The baseline at 0 h served as control to identify ASVs with an increased abundance compared to the beginning of the experiment. ASVs significantly enriched in both analyses were considered responsive to the amendment. For the mouse gut content microcosm experiments, each treatment was evaluated against the unamended control at each time point. In the conventional mouse experiment, ASV abundance was compared to the baseline at 0 h for each treatment. However, ASVs were considered significantly responsive to the amendment only, if they were not also significantly enriched in the unamended control over time. Adjusted *p*-values of <0.01 were considered significant. The taxonomic classification of enriched ASVs in all experiments was verified by Blastn against the NCBI 16S rRNA sequences database, excluding uncultured and environmental sample sequences [46].

The total 16S rRNA gene copies were quantified by real-time PCR (CFX96 Real-Time System; Bio-Rad Laboratories Ges.m.b.H, Austria). PCR reactions consisted of 10 μl iQ SyBr Green Mix 2x, 0.8 μl of each primer (0.4 µM), 6.4 μl sterile nuclease-free water, and 2 μl DNA extract. The universal primers 515F (5‘-GTG YCA GCM GCC GCG GTA A-3‘) [47] and 806R (5‘-GGA CTA CNV GGG TWT CTA AT-3‘) [48] targeting bacteria and archaea were used. The PCR program started with an initial denaturation step at 95°C for 5 min followed by 40 cycles of 95°C for 30 s, 55°C for 30 s, and 72°C for 30 s. Final elongation was performed at 72°C for 2 min. A negative control, i.e. water as template, was included in each run. Genomic DNA of *Escherichia coli* K12 was used to prepare a standard serial dilution series of 10^9^ (15.4 ng/µl DNA of the standard) to 10^3^ copy numbers of the 16S rRNA gene to generate a standard curve, which was linear across the entire dilution series. The 16S rRNA gene copy numbers of selected ASVs were calculated by multiplying their relative abundances with the total 16S rRNA gene abundances determined by real-time PCR.

### Analysis of genomes from the mouse microbiome

High-quality mouse microbiota-derived bacterial genomes were downloaded from the Mouse Gastrointestinal Bacteria Catalogue (n=26640) [49]. Open reading frames were determined using prodigal and subsequently annotated via HMSS2 [50]. Additional hidden Markov models were generated for SQDG and DHPS conversion related proteins not represented in HMSS2, as described previously [51]. Among others, this included isethionate reductase IsrBHDK [52], DHPS-sulfolyase HpsGH, DHPS-dehydratase HpfGH [17], sulfopropionaldehyde reductase HpfD, succinate semialdehyde dehydrogenase Ssadh [53], DHPS dehydrogenase DhpA, 3-sulfolactaldehyde dehydrogenase SlaB [16] and sulfoquinovosidase SqgA [7] (Table S1). For all hidden Markov models, a general bitscore cutoff of 100 was chosen to detect homologs. All hits were tested for co-location in the respective genome with a maximum distance of 3500 nucleotides between genes, as described before [51]. The resulting gene clusters were then searched for patterns of genes with the same sequential occurrence using a custom implementation of the colinear syntenic blocks finder algorithm [54], with a tolerance of two gene insertions and a length of four co-located genes. This was done to ensure enzyme function in the respective metabolism by sequence similarity and genetic vicinity.

## Results

### SQ is not degraded by human HT-29 cells *in vitro*

To test whether SQ is degraded by human intestinal cells, we incubated HT-29 human colorectal cancer cells with and without SQ. SQ concentrations in cell culture supernatants from both SQ-supplemented DMEM and glucose-free DMEM remained unchanged throughout the experiment (Figure 2 a, b). Lactate was only produced in glucose-containing incubations (Figure 2 c, d). The effect of increasing glucose or SQ concentrations on HT-29 cell growth in DMEM was similar for both substrates. Protein production was fully inhibited only in cultures treated with very high, non-physiological concentrations (250 mM) of glucose or SQ (Figure 2 e, f). Overall, these findings suggested that SQ is not degraded and metabolized by human cells *in vitro*. A potential limitation of using cancer cells is that their metabolism does not necessarily reflect the metabolism of primary cells [55].

**Figure 2:**
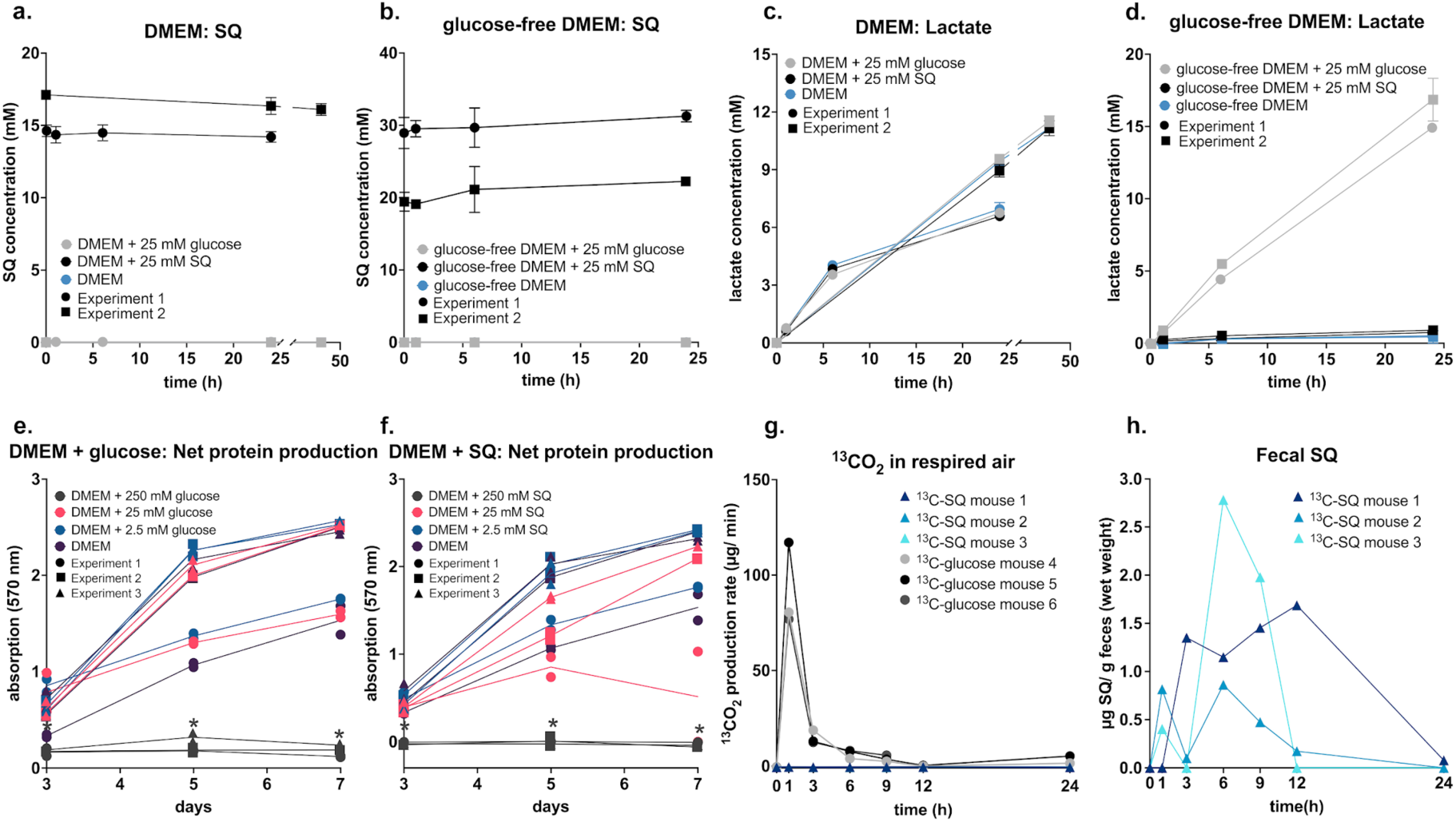
SQ was not metabolized by human intestinal cells and germ-free mice. SQ was not degraded by HT-29 cells in DMEM medium amended with 25 mM SQ (shown in black) (a). SQ was not degraded by HT-29 cells in glucose-free DMEM medium amended with 25 mM SQ (shown in black) (b). HT-29 cells produced lactate independent of the treatment in DMEM medium (c). In glucose-free DMEM, HT-29 cells produced lactate only in incubations amended with 25 mM glucose (shown in grey) (d). Protein production was inhibited in incubations with 250 mM glucose (shown in dark grey), but not in incubations with 25 mM glucose (shown in pink) and 2.5 mM glucose (shown in blue) (e). Protein production was inhibited in incubations with 250 mM SQ (shown in dark grey), but not in incubations with 25 mM SQ (shown in pink) and 2.5 mM SQ (shown in blue) (f). Each germ-free mouse (n = 3 per group) was gavaged with 10 mg of fully labeled ^13^C-glucose or ^13^C-SQ. (g). Glucose, but not SQ, was respired by germ-free mice as shown by ^13^CO2 production via breath air isotope analysis. SQ was repeatedly detected in the feces of all mice gavaged with ^13^C-SQ within 24 h after the gavage (h). In a., b., c., and d., lines show averages of triplicate incubations with error bars representing one standard deviation. Two experiments were conducted varying only slightly in the sampling time points. Different data point shapes indicate the two experiments. In e. and f., lines show averages of duplicate measurements and data points show individual values. Different data point shapes represent the three conducted experiments. Asterisks mark significant differences between a treatment group and the control (DMEM medium without amendment) at each time point. Significant differences were determined by ANOVA with Dunnett’s post hoc test. Adjusted *p*-values smaller than 0.05 were regarded as significant. SQ, sulfoquinovose; DMEM, Dulbecco’s Modified Eagle Medium.

### 13C-labeled SQ is not respired to ^13^C-CO_2_ by germ-free mice

Next, we employed stable isotope tracing with ^13^C-SQ in germ-free mice to study potential SQ degradation by mammalian cells *in vivo*. Germ-free mice received a single dose (10 mg) of either fully labeled ^13^C-SQ or ^13^C-glucose and their respired air and fecal samples were subsequently collected over a 24-hour period for ^13^CO_2_ breath tests and metabolite analysis, respectively. Mice administered ^13^C-SQ did not produce ^13^CO_2_. Contrarily, mice receiving ^13^C-glucose produced up to 117 µg ^13^CO_2_/min at one hour post-gavage (Figure 2 g). In mice gavaged with ^13^C-SQ, SQ was detected in fecal samples already one hour after its administration (Figure 2 h). Yet, SQ degradation metabolites were not detected in the feces.

### Response of the microbiota to different doses of SQ in human fecal microcosms

We performed anoxic incubations with human fecal microcosms to assess the dose-dependent effects of SQ (0.01 mM, 0.1 mM, 1 mM, and 10 mM) on metabolite production and gut microbiota composition (Figure 3 a). Only the microcosms with high SQ concentrations (1 mM, 10 mM) consistently showed significant differences in metabolite levels compared to the unamended control (Figure S1). The temporal dynamics of SQ degradation and the production of metabolic products were consistent between the 1 mM and 10 mM SQ treatments (Figure 3 b, c). SQ was mostly degraded within 12 h, coinciding with the production of DHPS. In comparison, SL and sulfide were produced later and represented only a smaller fraction of the produced sulfur metabolites within the 24 h incubation. SQ, DHPS, and SL were not detected in the unamended control incubation (Figure 3 d). The production of acetate and butyrate, known SQ degradation products [15,16,39], increased significantly in the 1 mM and 10 mM SQ treatments compared to the unamended control (Figure 3 e, f).

**Figure 3:**
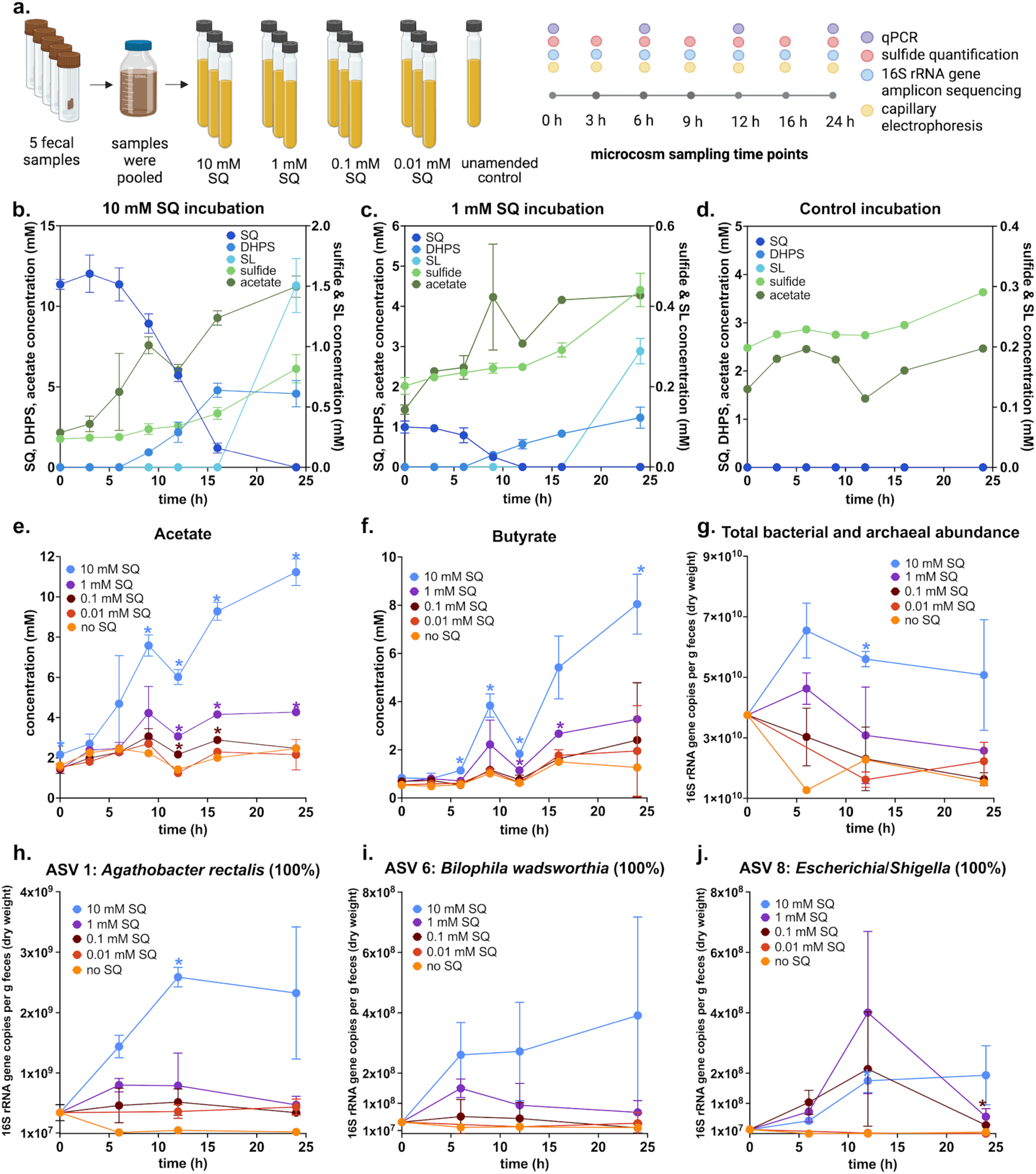
Metabolite dynamics and differential responses of microbiota members to different doses of SQ in 24 h human fecal microcosm incubations. Overview of the experimental and analytical approach (created with biorender.com) (a). Sulfur metabolite dynamics in incubations amended with 10 mM SQ (b), 1 mM SQ (c), and in the unamended control (d). SQ, DHPS, and SL were not detected in unamended control microcosms. Higher doses of SQ led to increased concentrations of acetate (e) and butyrate (f). No significant differences in acetate concentrations were detected between incubations with 0.01 mM SQ (shown in red) and unamended controls (shown in orange). No significant differences in butyrate concentrations were detected between incubations with 0.1 mM SQ (shown in dark red) or 0.01 mM SQ (shown in red) and unamended controls (shown in orange). Changes in absolute bacterial and archaeal 16S rRNA gene copy numbers in the microcosm incubations with different SQ doses (g). Changes in the absolute abundance of *A. rectalis* ASV 1 (h), *B. wadsworthia* ASV 6 (i), and *Escherichia/Shigella* ASV 8 (j). The 16S rRNA gene sequence similarity (%) to the closest, named relative is shown for each ASV. In b., c., e., f., g., h., i., and j., the lines show the average of triplicate incubations with error bars representing one standard deviation. Only a single incubation was performed for the unamended control microcosm. Coloured asterisks show significant differences between the respective treatment (n = 3) and the retrospectively designated control incubations (n = 4, consisting of three 0.01 mM SQ incubations and one unamended control) at individual time points. Significant differences were determined by ANOVA with Dunnett’s post hoc test. Adjusted *p*-values smaller than 0.05 were regarded as significant. SQ, sulfoquinovose; DHPS, 2,3-dihydroxypropane-1-sulfonate; SL, 3-sulfolactate; ASV, amplicon sequencing variant.

Changes in total microbial abundance during the 24 h incubation period were quantified by 16S rRNA gene-targeted real-time PCR (Figure 3 g). Gene copy numbers decreased in microcosms without SQ and with 0.01 or 0.1 mM SQ and remained transiently constant in microcosms with 1 mM SQ. A significant increase in gene copy numbers was only observed in microcosms with 10 mM SQ.

We performed 16S rRNA gene amplicon sequencing and differential abundance analyses to identify amplicon sequence variants (ASVs) with increased relative abundance in SQ-amended incubations and thus a possible role in SQ degradation. Significantly enriched ASVs were only detected in incubations with concentrations of 0.1 mM SQ or higher; one, two, and fourteen ASVs were enriched in microcosms with 0.1, 1, and 10 mM SQ, respectively (Table S2, Figure S2). *Escherichia/Shigella* ASV 8 was enriched at all three SQ concentrations, *Waltera intestinalis* ASV 7 was enriched at 1 and 10 mM SQ, while the following ASVs were only enriched at 10 mM SQ: *A. rectalis* ASV 1, *Alistipes shahii* ASV 2, *Ruminococcus bromii* ASV 3, *Oscillospiraceae* ASV 5, *B. wadsworthia* ASV 6, *Enterocloster citroniae* ASV 9, *Phocaeicola dorei* ASV 4, *Phocaeicola massiliensis* ASV 12, *Phocaeicola vulgatus* ASV 14, *Bacteroides xylanisolvens* ASV 10, *Bacteroides caccae* ASV 11, and *Bacteroides ovatus* ASV 13. Variations in the dose-response patterns of SQ-responsive microorganisms are not yet fully explained but may result from differences in SQ metabolic energetics and enzyme kinetics. Close relatives of some of the enriched ASVs (Figure 3 h, i, j) have been physiologically confirmed to degrade SQ (*A. rectalis*, *E. coli*) or DHPS (*B. wadsworthia*) [10,15]. Furthermore, members of the genera *Enterocloster* and *Ruminococcus* encode the sulfofructose-transaldolase pathway [15]. Together with a member of *Alistipes*, members of these genera/species have also previously been significantly enriched in human fecal microcosms amended with 10 mM SQ [15]. ASVs related to these taxa generally showed a continuous increase in absolute abundance over the 24 h incubation, which coincided with the kinetics of SQ disappearance and DHPS production (Figures 3 and S2). In contrast, ASVs of *Phocaeicola* and *Bacteroides* showed an initial spike in absolute abundances at 6 h followed by a decline, suggesting they are not directly involved in SQ degradation (Table S2, Figure S2). Type strain genomes of the most closely related *Alistipes*, *Waltera*, *Bacteroides* or *Phocaeicola* species do not encode known SQ or DHPS degradation pathways.

### Incomplete SQ degradation in mouse gut content microcosms

Two independent microcosm experiments were performed with pooled gut contents of mice and amended with SQ or DHPS to investigate the SQ metabolism (Figure 4 a). In experiment 1, microcosms containing entire gut contents were incubated with either 10 mM SQ or 10 mM DHPS. Following a lag phase of over 24 hours, SQ degradation to DHPS was observed in SQ-amended incubations at a near 1:1 ratio, as measured by liquid chromatography-mass spectrometry (Figure 4 b). SL, sulfoacetate, and 3-HPS were not detected. Yet, small amounts of isethionate (0.01 mM after 48 h and 0.02 mM after 96 h) were detected (Figure S3). Sulfide concentrations in SQ-amended microcosms increased slightly, remaining below 1 mM, and did not correlate with SQ degradation in timing or stoichiometry. Capillary electrophoresis analysis of SCFAs showed an increase in acetate production in SQ-amended incubations, from 5 mM to 11.1 mM over 96 h, compared to the control incubation, from 2.6 mM to 4.3 mM (Figure 4 b). Concentrations of DHPS and acetate remained constant in DHPS-amended incubations (Figure 4 c). Sulfide concentrations were significantly higher in DHPS-amended incubations compared to SQ-amended incubations and unamended controls. However, the small increase from 0.2 mM to 0.5-0.6 mM sulfide in the DHPS-amended incubations compared to 0.3-0.4 mM sulfide in the unamended control did not suggest a quantitative turnover of DHPS to sulfide (Figure 4 c). SQ and DHPS were not detected in the unamended control incubations, and sulfide concentrations remained below 0.4 mM (Figure 4 d).

**Figure 4:**
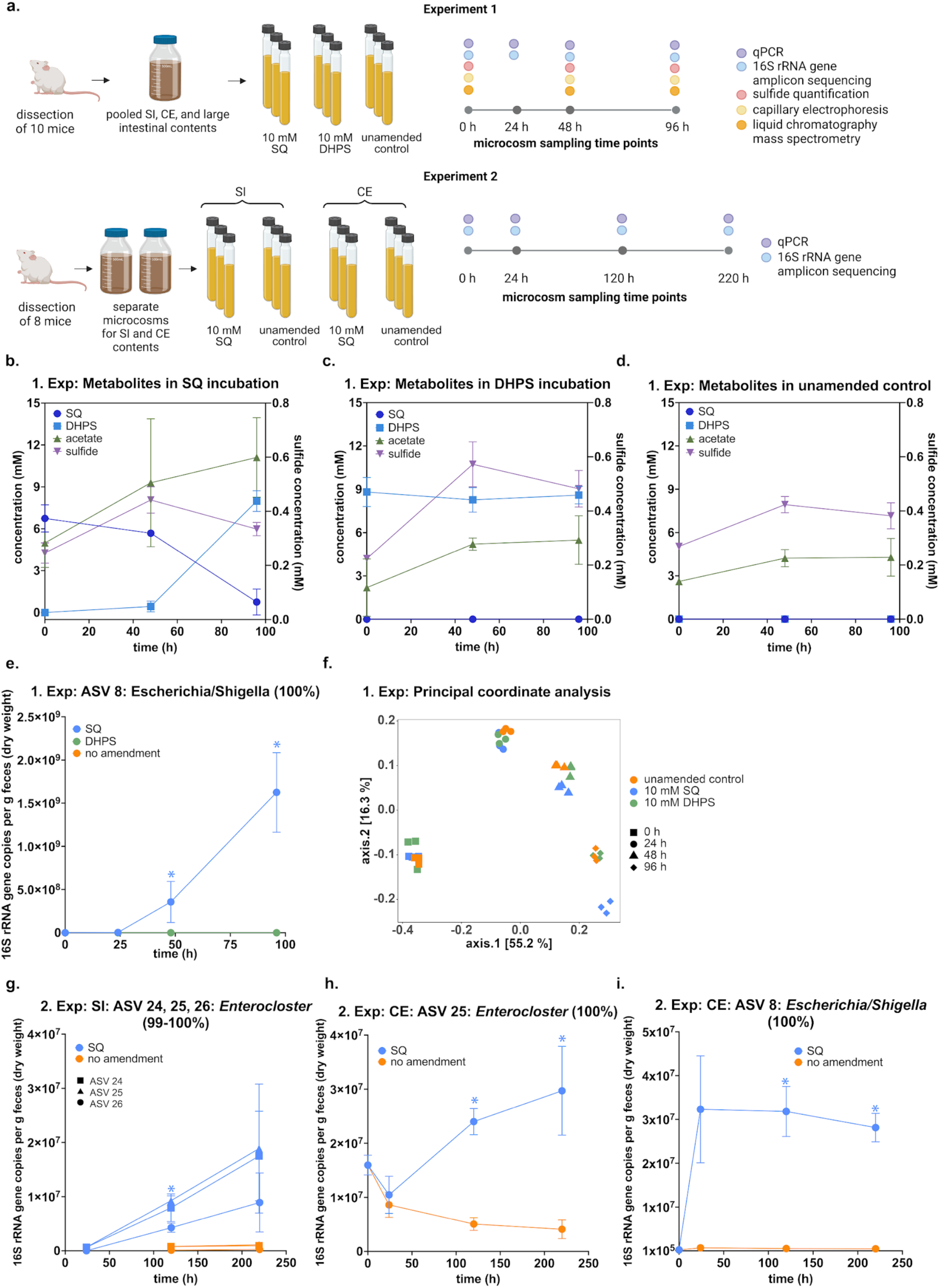
Metabolite dynamics and differential responses of microbiota showed that SQ, but not DHPS, was degraded in murine gut content microcosms. Overview of the experimental and analytical approach (created with biorender.com) (a). Metabolite dynamics in gut content microcosms (Experiment 1: pooled small intestinal, cecal, and large intestinal contents) incubated with 10 mM SQ (b), 10 mM DHPS (c), and without amendment (control) (d). SQ and DHPS were not detected in control incubations. Calculated absolute 16S rRNA gene abundance of *Escherichia*/*Shigella* ASV 8 in incubations of experiment 1 (e). Principal coordinate analysis plot of beta diversity (Bray-Curtis dissimilarity) of 16S rRNA gene amplicon data from incubations of experiment 1 (f). Calculated absolute 16S rRNA gene abundance of selected ASVs that were significantly enriched in SQ-amended mouse small intestinal content and cecum content incubations of experiment 2 (g-i). The 16S rRNA gene sequence similarity (%) to the closest, named relative is shown for each ASV. The lines in b., c., d., e., g., h. and i. show averages of triplicate incubations with error bars representing one standard deviation. Coloured asterisks show significant differences between the respective treatment and the unamended control at individual time points. Significant differences were either determined by two-way ANOVA with Dunnett post hoc test (1. experiment: pooled gut content) or by multiple unpaired t-tests with Holm-Šidák post hoc test (2. experiment: small intestine and cecum). Adjusted *p*-values smaller than 0.05 were regarded as significant. SQ, sulfoquinovose; DHPS, 2,3-dihydroxypropane-1-sulfonate; Exp, experiment; SI, small intestine; CE, cecum; ASV, amplicon sequencing variant.

Microbial abundances decreased over time in all microcosms, but remained significantly higher in SQ incubations from 48 h onwards (Figure S4). Microbial abundances decreased from 1.8 ± 0.7 x 10^10^ 16S rRNA gene copies/g dry weight to 5.2 ± 1.0 x 10⁹ 16S rRNA gene copies/g dry weight in SQ-amended microcosms and to 5.3 ± 1.4 x 10⁸ 16S rRNA gene copies/g dry weight in the unamended controls.

Differential abundance analysis of 16S rRNA gene amplicon sequencing data identified nine ASVs enriched in the 10 mM SQ incubations compared to unamended controls (Figure S5). *Escherichia/Shigella* ASV 8 had the highest log_2_ fold change of all enriched ASVs and its absolute abundance dynamic corresponded to the turnover of SQ into DHPS (Figure 4 b, e). Other enriched ASVs included *Lachnospiraceae* ASV 15, *Bacteroides acidifaciens* ASV 16, *Bacteroides* ASV 19, *F. rodentium* ASV 17, *Coprobacillaceae* ASV 18, *Ruminococcus* ASV 20, *Prevotellaceae* ASV 21, and *P. vulgatus* ASV 14 (Table S3). Notably, *Escherichia/Shigella* ASV 8 and *P. vulgatus* ASV 14 were enriched in both human fecal microcosms and mouse gut content microcosms with SQ. In contrast to SQ-amended incubations, temporal changes in microbial community composition in the DHPS-amended incubations were indistinguishable from control incubations and no ASVs were identified as enriched (Figure 4 f).

For experiment 2, separate microcosms for small intestinal contents and cecal contents were each incubated with 10 mM SQ and analyzed for changes in microbial community composition compared to unamended control incubations. Microbial 16S rRNA gene copy numbers decreased in control incubations, while remaining rather constant in cecal incubations with SQ or decreased less in small intestinal incubations with SQ compared to the control (Figure S4). Seven ASVs were significantly enriched in small intestinal incubations and four ASVs in cecal incubations. ASVs in small intestinal incubations were *Enterococcus* ASV 22 and 23, *Enterocloster* ASV 24, 25, and 26, *Ligilactobacillus* ASV 27, and *Akkermansia muciniphila* ASV 28 (Figure 4 g, Table S3). In cecal incubations, enriched ASVs belonged to *Escherichia/Shigella* ASV 8, *Parabacteroides distasonis* ASV 29 and 30 and, *Enterocloster* ASV 25 (Figure 4 h, i, Table S3). In contrast to the other ASVs, the absolute abundances of *Ligilactobacillus* ASV 27 and *A. muciniphila* ASV 28 decreased during the incubation and thus are unlikely to be involved in primary SQ degradation (Figure S6).

### SQ is degraded to DHPS in conventional wild type mice

To investigate SQ metabolism by the mouse gut microbiota *in vivo*, we gavaged a single dose of either 1 mg SQ or 10 mg SQ to conventional wild type laboratory mice and collected fecal samples up to 24 h post-gavage (Figure 5 a). Mice fed with SQ showed no signs of digestive problems or general discomfort. While the water content of fecal pellets was not measured, the consistency of fecal pellets remained stable throughout the experiment, with no visible difference between the tap water control and SQ-treated mice. Fecal metabolites were analyzed via capillary electrophoresis as well as liquid chromatography-mass spectrometry (Table S4). SQ was detected in mice receiving 10 mg SQ between 3 and 6 h post-gavage, while DHPS appeared at 6 h and remained detectable in some mice until the end of the experiment (Figure 5 b, Figure S7). Additionally, taurine as well as small amounts of 3-sulfolactate, sulfoacetate, and isethionate were measured via liquid chromatography-mass spectrometry in mice receiving 10 mg SQ (Table S4). Taurine and isethionate were detected already at the beginning of the experiment, indicating that the presence of these metabolites was independent from SQ addition. Temporal dynamics of SCFA concentrations, total 16S rRNA gene copy numbers, and microbial composition were not significantly different between the three treatment groups (Figure 5 c - g).

**Figure 5:**
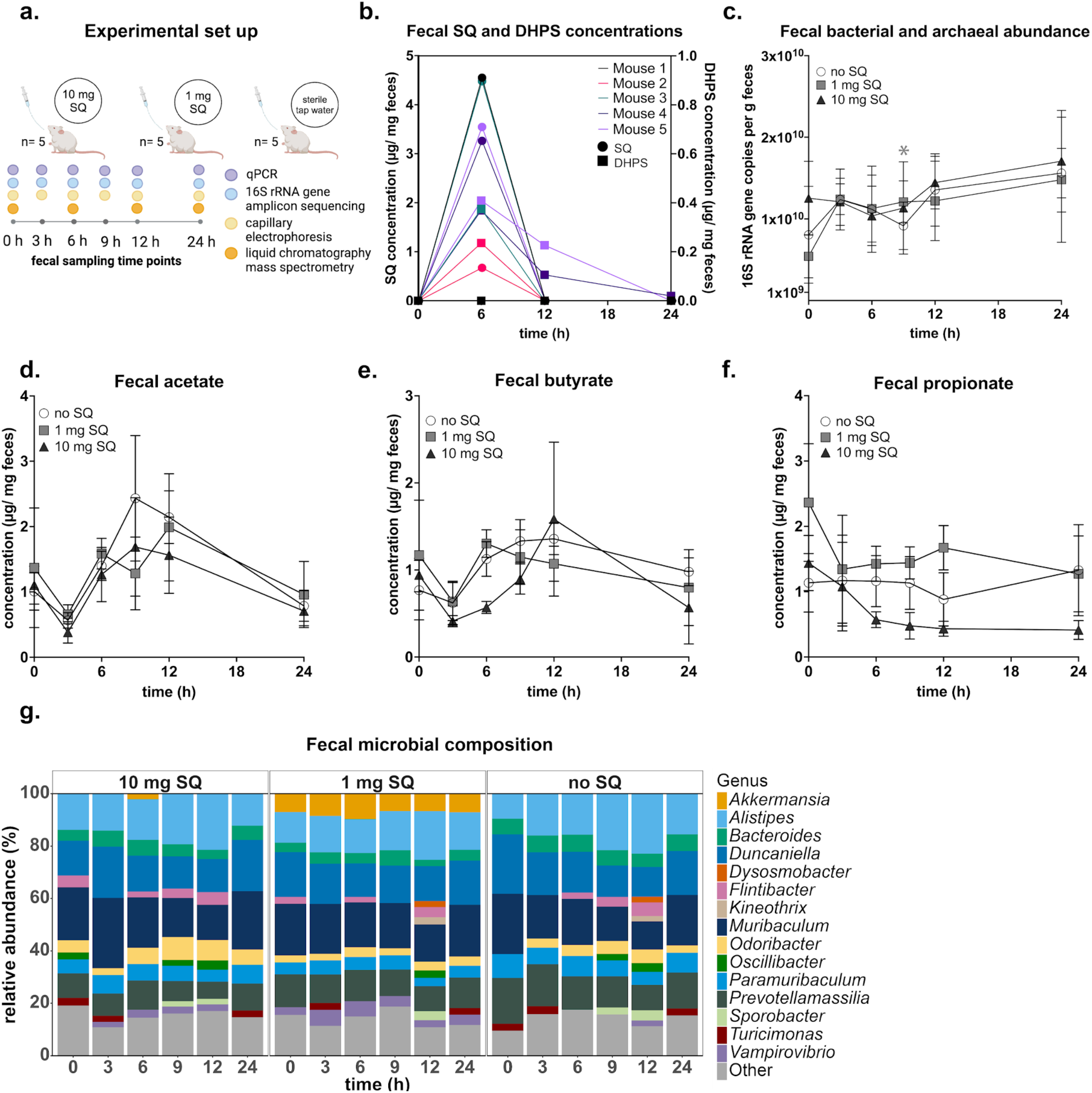
Fecal metabolite and microbial community dynamics of conventional wild type mice over 24 h after receiving a single dose of SQ. Overview of the experimental and analytical approach (created with biorender.com) (a). Concentrations of SQ (circles) and DHPS (boxes), as analyzed by liquid chromatography-mass spectrometry, in feces of mice (n = 5) that each received a single dose of 10 mg SQ by gavage (b). Metabolite concentrations of all treatment groups, as analyzed by capillary electrophoresis, are depicted in Supplementary Table S4 and Figure S7. Total bacterial and archaeal 16S rRNA gene copy numbers in feces of the three treatment groups (c). Five biological replicates were measured in triplicate qPCR assays. The line shows the mean and the error bars represent one standard deviation. Concentrations of acetate (d), butyrate (e), and propionate (f) in feces of the three treatment groups. Mean values of five biological replicates are shown and error bars depict one standard deviation. Mean relative 16S rRNA gene abundance of dominant bacterial genera in feces of the three treatment groups (g). All genera below an abundance of 2% are summarized in ‘Other’. All data refer to the wet weight of the feces. Asterisks show significant differences between the respective treatment and the tap water control at individual time points. Significant differences in c., d., e., and f. were determined by ANOVA with Dunnett’s post hoc test. Adjusted *p*-values smaller than 0.05 were regarded as significant. SQ sulfoquinovose, DHPS 2,3-dihydroxypropane-1-sulfonate.

### SQDG/SQ and DHPS degradation pathways are encoded in few bacterial genomes from the mouse gut

We comprehensively examined the genetic potential for SQDG/SQ and DHPS degradation in the mouse microbiota by analyzing the collection of 26640 metagenome-assembled genomes (MAGs) from the Mouse Gastrointestinal Bacteria Catalogue for presence of the respective pathways [49]. Our analysis revealed SQDG/SQ and DHPS degradation pathways in only 0.3% (n=76) and 0.2 % (n=48) of all MAGs and in 16 (1.5%) and 8 (0.7%) of the 1094 species-level genome bins, respectively (Figure 6 a, b). The sulfofructose-transaldolase pathway was encoded in 57 MAGs, primarily from *E. clostridioformis*, *Otoolea symbiosa* (formerly *Clostridium symbiosum*), and *E. aldenensis*, and represented the most prevalent SQDG/SQ degradation pathway in the mouse microbiome (Figure 6 a). The sulfo-Embden-Meyerhof-Parnas pathway was encoded in 13 *Enterobacteriaceae* MAGs. The sulfo-transketolase pathway was encoded in 6 *Oscillibacter* MAGs. Key enzymes in DHPS degradation pathways (Figure 1 b) were encoded in 48 MAGs across five genera. These included 16 MAGs with *hpsN*, 12 MAGs with *dhpA*, 10 MAGs with *hpfGH*, 8 MAGs with *hpsGH*, and 2 MAGs with both *hpsGH* and *dhpA* (Figure 6 b). These pathways could generate sulfite intracellularly and were present in the sulfidogenic genera *Desulfovibrio*, *Taurinivorans*, *Mailhella*, and *Bilophila* that employ a DsrAB-dissimilatory sulfite reductase to respire sulfite to H_2_S [15,16,19]. Nine *E. clostridioformis* MAGs and one *E. aldenensis* MAG encoded the fermentative *hpfGHD* pathway along with the sulfofructose-transaldolase pathway, suggesting these bacteria may ferment SQ via DHPS also to 3-HPS [17].

**Figure 6:**
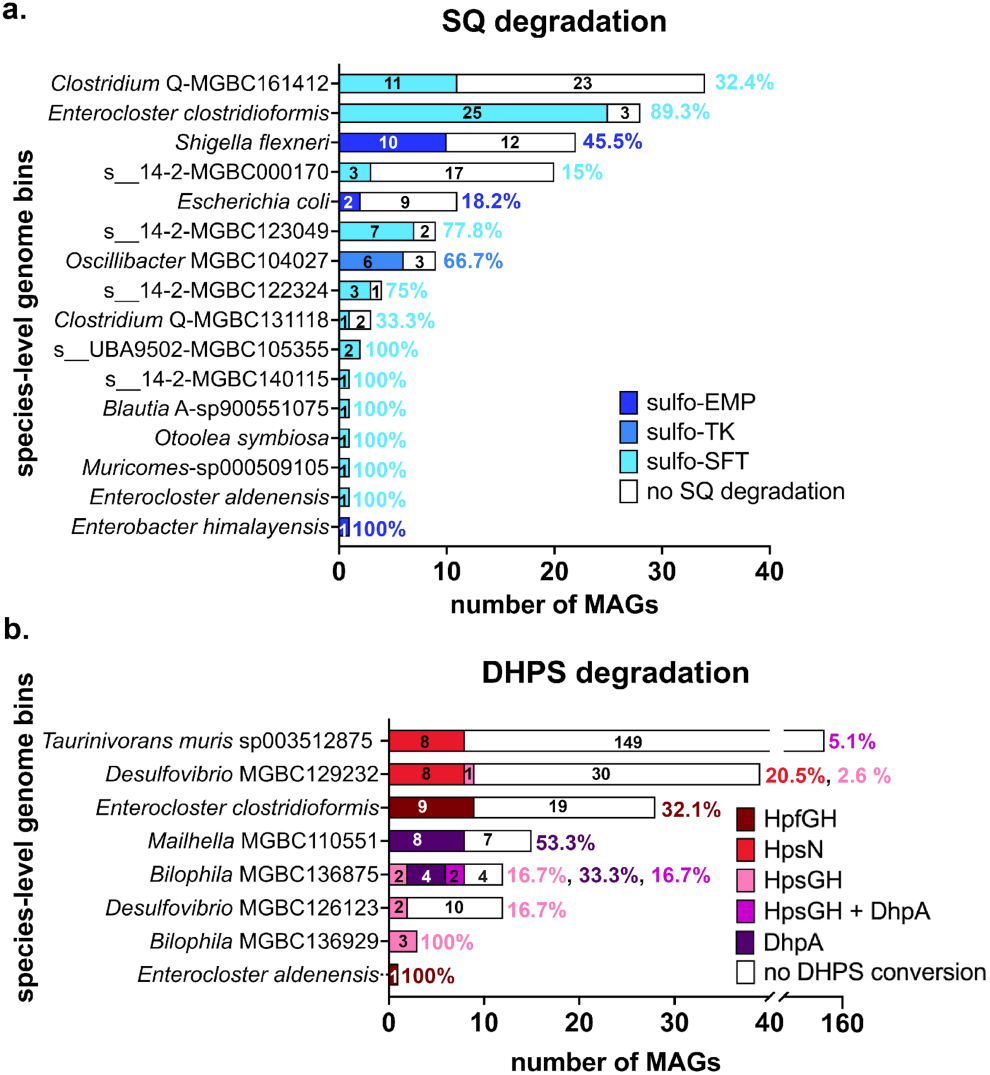
Analysis of microbial genomes revealed genetic SQDG/SQ and DHPS degradation potential is sparsely distributed in the mouse microbiome. Distribution of SQDG/SQ (a) and DHPS (b) degradation potential across a collection of 26,640 microbial MAGs from the mouse intestinal tract. Species-level genome bins are ordered based on the total number of genomes. The total and relative (%) number of genomes that encode a specific SQDG/SQ or DHPS pathway (shown in different colors) is indicated for each species-level genome bin. Sixteen (represented by 76 MAGs) and eight (represented by 48 MAGs) species-level genome bins, contained complete gene clusters for SQDG/SQ and DHPS degradation, respectively. Some MAGs of *E. clostridioformis* and *E. aldenensis* encoded both SQDG/SQ and DHPS degradation pathways. MAG, metagenome-assembled genome; sulfo-EMP, sulfoglycolytic Embden-Meyerhof-Parnas pathway; sulfo-ED, sulfoglycolytic Entner–Doudoroff pathway; sulfo-SFT, sulfoglycolytic sulfofructose-transaldolase pathway; SQ, sulfoquinovose; DHPS, 2,3-dihydroxypropane-1-sulfonate; HpsG, DHPS-sulfolyase; HpsH, DHPS-sulfolyase activating enzyme; HpfG, DHPS-dehydratase; HpfH, DHPS-dehydratase activating enzyme; HpsN, DHPS-1-dehydrogenase; DhpA, dehydrogenase.

### *Enterocloster clostridioformis* degrades SQ to DHPS in pure culture

Our mouse gut content microcosm experiments and genome analyses revealed members of the genus *Enterocloster* as SQ degraders. We thus performed anaerobic growth tests with the mouse strain *E. clostridioformis* YL32, which shares 100%, 99.6%, and 99.2% 16S rRNA gene sequence identity with the enriched ASVs 25, 24, and 26 in the mouse gut microcosms, respectively. Initial growth of strain YL32 in diluted or undiluted rich medium did not differ, regardless of SQ supplementation. However, cultures supplemented with SQ exhibited a secondary growth phase during the late stationary phase, achieving significantly higher optical densities than cultures without SQ. This secondary growth was more pronounced in diluted rich medium (Figure 7 a). Diauxic growth in SQ-supplemented cultures indicated that SQ supported growth only after energetically more favorable substrates in the rich medium were consumed. This was supported by the timing of SQ degradation, which concurred with the second growth phase after 100 h. SQ was degraded nearly stoichiometrically to DHPS (Figure 7 b) while sulfide production did not differ significantly between cultures with and without SQ (Figure 7 c). Although *E. clostridioformis* YL32 possesses the genetic potential to ferment DHPS further (Figure 7 d), the DHPS fermentation product, 3-HPS, was not detected.

**Figure 7:**
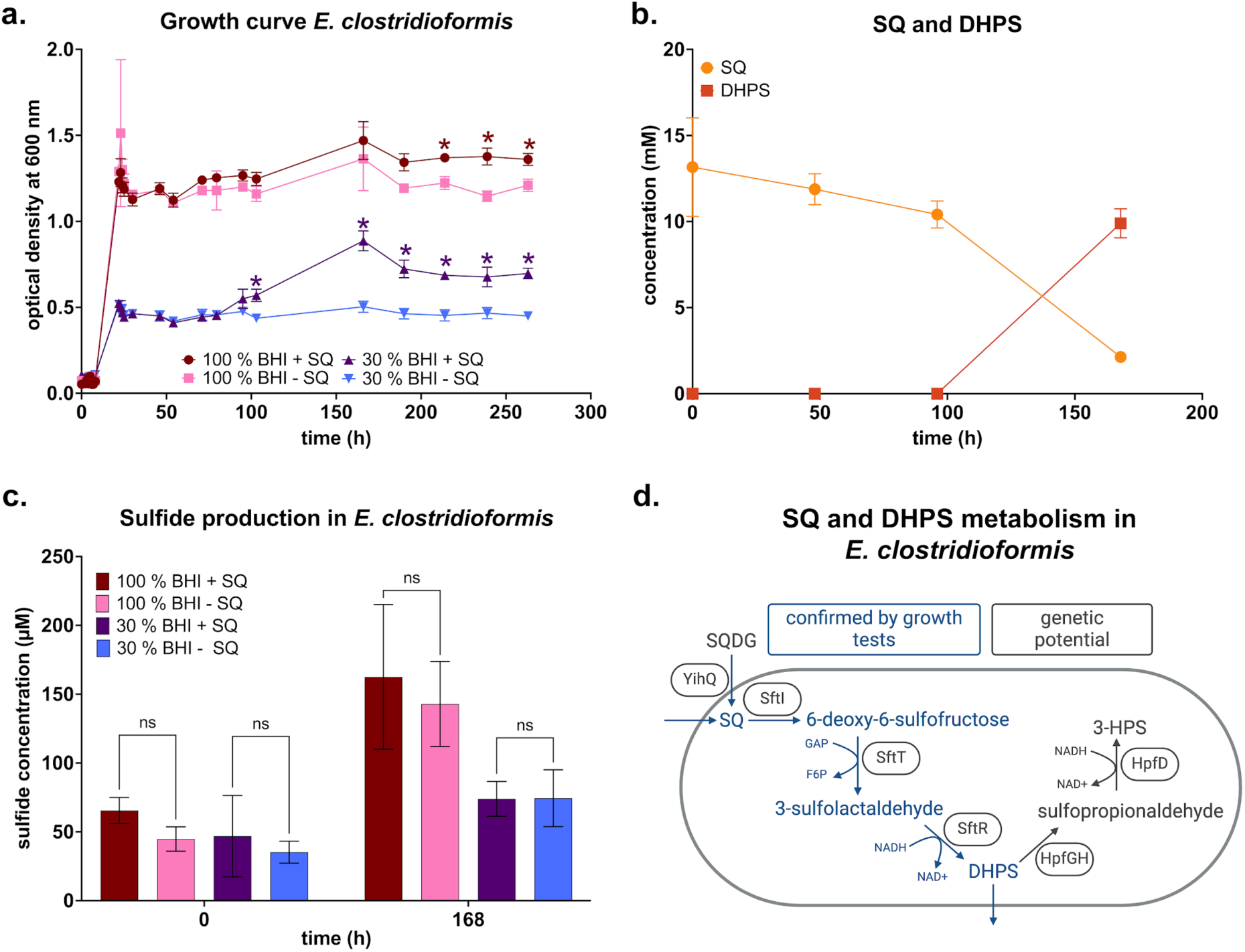
A pure culture of *Enterocloster clostridioformis* anaerobically degraded SQ to DHPS. Anaerobic growth tests of *E. clostridioformis* strain YL32 with 10 mM SQ (a). SQ triggered a second growth phase of *E. clostridioformis* in diluted (30%) rich medium (a) that coincided with a decrease of SQ and an increase of DHPS at a stoichiometry of about 1:1 (b). Small amounts of sulfide were produced in all treatments but there were no significant differences between the sulfide produced in incubations amended with SQ compared to unamended control incubations (c). Schematic overview of the proposed SQ and DHPS pathways of *E. clostridioformis* YL32 (genome assembly number GCA_016696785) based on the growth tests and genome analysis (created with biorender.com) (d). SQ can either be imported via the transporter SftA or cleaved from SQDG by the sulfoquinovosidase YihQ (gene locus tag I5Q83_09300). SQ is converted to 6-deoxy-6-sulfofructose via the SQ isomerase SftI (gene locus tag I5Q83_09270). The 6-deoxy-6-sulfofructose transaldolase SftT (gene locus tag I5Q83_09315) catalyzes the transfer of a C3-moiety from 6-deoxy-6-sulfofructose to glyceraldehyde-3-phosphate (GAP), forming fructose 6-phosphate (F6P). The resulting 3-sulfolactaldehyde is then reduced to DHPS by the 3-sulfolactaldehyde reductase SftR (gene locus tag I5Q83_09310). The DHPS dehydratase HpfG (gene locus tag I5Q83_10645) and its activating enzyme HpfH (gene locus tag I5Q83_10640) catalyze the cleavage of the CO-bond of DHPS to form sulfopropionaldehyde, which is then reduced by the sulfopropionaldehyde reductase HpfD (gene locus tag I5Q83_10625) to 3-HPS. In a., b., c., lines/bars show averages of triplicate incubations with error bars representing one standard deviation. In a., coloured asterisks show significant differences between treatment groups and their respective control groups without SQ. In c., brackets show non-significant results in pairwise comparisons of treatment groups and their respective unamended control. Significance was tested using ANOVA with Tukey’s post hoc test. Adjusted *p*-values smaller than 0.05 were regarded as significant. SQ, sulfoquinovose; BHI, brain-heart infusion medium; DHPS, 2,3-dihydroxypropane-1-sulfonate; 3-HPS, 3-hydroxypropane-1-sulfonate; ns, not significant; SQDG, sulfoquinovosyldiacylglycerol.

Additionally, we tested SQ degradation by the mouse strain *F. rodentium* ALO17, which shares 100% identity with an ASV enriched in the mouse gut microcosms. However, growth of strain ALO17 was not significantly different between cultures with and without SQ. Also, SQ degradation and formation of degradation products were not detected (Figure S8). These results are in line with the absence of known SQ and DHPS pathway genes in the *F. rodentium* ALO17 genome.

## Discussion

SQ in the gut exclusively derives from dietary intake, primarily of green edible plants (e.g. spinach, green onions) and edible micro- and macroalgae (e.g. spirulina, nori) that contain the ubiquitous thylakoid membrane lipid SQDG. In the mammalian gut, pancreatic lipases hydrolyze SQDG to sulfoquinovosyl glycerol [56,57]. Subsequently, SQ is cleaved off from SQDG or sulfoquinovosyl glycerol by sulfoquinovosidases [6,7], key enzymes of certain gut bacteria that take up SQ for further respiration or fermentation. SQ is structurally analogous to glucose-6-phosphate and was initially proposed to be degraded via a pathway similar to glycolysis [58,59]. Since the first discovery of sulfoglycolysis in bacteria [10], further sulfoglycolytic pathways have been described in different bacterial taxa [8,9,11–14]. While plants [58] and algae [60] can also degrade SQ via yet unknown pathways, it was unclear if animals are capable of SQ degradation and if mammalian cells are involved in intestinal SQ metabolism. Here, we show that SQ is neither metabolized by human colorectal adenocarcinoma (HT-29) cells *in vitro* nor respired to CO_2_ by germ-free mice *in vivo*. This suggests that SQ cannot be metabolized by mammals, and instead serves exclusively as a nutrient and growth substrate for specialized bacteria in the animal gut.

A. *rectalis* was previously shown to be the most dominant SQ-degrading species in human fecal samples. Other species, including *F. prausnitzii*, *E. aldenensis* (formerly *Clostridium aldenense*), *E. clostridioformis* (formerly *Clostridium clostridioforme*), and *Roseburia* sp. AM16-25 also contributed to transcription of sulfofructose-transaldolase pathway genes in the fecal microbiome [15]. Transcription of the sulfo-Embden-Meyerhof-Parnas pathway of *Enterobacteriaceae* such as *E. coli* was considerably less important. Additionally, *B. wadsworthia* was identified as the dominant degrader of DHPS, which is produced by the primary SQ fermenters [15]. Our human fecal microcosm experiments with different SQ doses corroborated the relevance of *A. rectalis* and *B. wadsworthia* for complete SQ degradation to H_2_S in the human gut. We further revealed that a minimum concentration of 0.1 mM SQ was sufficient to induce significant changes in the absolute abundance of selected microbiota members. The total amount of SQ in 10 ml microcosms amended with 0.1 mM SQ corresponds to the SQ content in about 11 g of fresh spinach (Table S5), the edible plant with the highest known SQDG concentration [3,61]. These results align with previous findings showing that SQ degradation ranks among the top third of all transcribed metabolic pathways of the fecal microbiome, establishing it as a microbial core function in the human gut [15]. Intestinal SQ levels reached by regular dietary intake of green vegetables may thus impact gut microbiota composition and potentially affect human health via SQ degradation metabolites [15]. The main SQ fermentation products are formate, acetate, and DHPS [10,13,16,62].

Besides acetate formation, our human fecal microcosm experiments also revealed a significantly higher butyrate concentration in microcosms amended with 10 mM SQ compared to the control (Figure 3 f). Increased butyrate production during SQ degradation has not been observed in previous SQ-amended human fecal microcosms [15], which suggests it may be specific to the mixture of human fecal samples used in our study. While the reason for the increased butyrate production remains unknown, it is likely produced by *A. rectalis* and other bacteria that were enriched in the SQ-amended microcosms. *A. rectalis* and relatives of other enriched ASVs (e.g. *W. intestinalis*) can produce butyrate from fermentation of carbohydrates [63–65]. Butyrate production by *A. rectalis* involves the acetyl-CoA pathway, with butyryl-CoA:acetate CoA-transferase (But) as the key terminal enzyme that generates butyrate from two acetyl-CoA molecules and acetate as co-substrate [66–69]. *Faecalicatena* sp. DSM22707, an isolate from petroleum reservoir produced water, produces butyrate and other metabolites from SQ, highlighting butyrate as a new SQ fermentation product [39]. Production of SCFAs, particularly of butyrate, is considered a beneficial function of the gut microbiota and a prominent effect of dietary supplementation with known prebiotics such as resistant starch, inulin-type fructans, and fructooligosaccharides [70–74]. Butyrate is the preferred energy source of the colonocytes [75] and has been linked to the prevention and treatment of colitis [76,77] and colorectal cancer [78]. Supporting the prebiotic potential of SQ, *A. rectalis* maintained stable abundances when switching from a high-fiber to a low-fiber diet supplemented with SQ in a humanized gnotobiotic mouse model colonized with fourteen fiber- and mucin-degrading bacterial strains from the human gut. Without SQ, its abundance dropped approximately 1000-fold after switching to the low-fiber diet. This effect was specific to *A. rectalis* and not observed for *O. symbiosa*, another SQ degrader in the consortium [79].

The identity and metabolic features of microorganisms involved in SQ degradation in the gut of non-human animals is unknown. Laboratory mice are important models of biomedicine and nutrition research. Here we demonstrate that, similar to the human gut, the capacity for SQ degradation is restricted to few members of the murine microbiota. For example, the sulfofructose-transaldolase pathway, the most prevalent SQ degradation pathway in both systems, is found in only 0.5% of human and 1.1% of mouse gut microbiota species-level genome bins, underscoring its limited distribution (Figure 6) [15]. SQ is mainly degraded via the sulfo-Embden-Meyerhof-Parnas pathway and the sulfofructose-transaldolase pathway to DHPS. Members of the genera *Escherichia* and *Enterocloster* are primary SQ degraders in the mouse gut. The more than 24 h lag phase till onset of SQ degradation in mouse gut content microcosms and the absence of a significant shift in fecal microbiota composition upon single-dose gavage of SQ to mice suggested that microbial SQ metabolism was not active before SQ supplementation. The SQDG contents of mouse chows are not known. However, the absence of SQ in feces before gavaging (Figures 2 h and 5 b) may indicate that the standard mouse chow used in this study, and possibly also in other mouse facilities, is not composed of foods with high SQDG content [3,80,81]. The single 1 mg and 10 mg SQ doses gavaged to mice correspond to the SQ content in approximately 0.37 g and 3.7 g of fresh spinach, which would represent about 7.5% and 75% of the total daily food intake in grams of a mouse, respectively (Table S5). Mice showed no adverse effects after ingesting a high SQ dose. Our results support previous findings from oral SQ supplementation experiments in susceptible IL-10-deficient mice [82]. Daily gavage with 0.45 mg SQ per g body weight, which corresponds roughly to the 10 mg SQ dose used in our study, over three weeks did not induce changes in microbial beta-diversity and intestinal inflammation in the IL-10-deficient mice [82].

Although some mouse microbiota members encode the capacity to either respire DHPS to H_2_S (*T. muris*, *Desulfovibrio* species) or ferment DHPS to 3-HPS (*E. clostridioformis*), experiments with gut content microcosms, with conventional laboratory mice, and mouse gut-derived bacterial strains did not provide evidence for active degradation of DHPS. *T. muris* did not utilize DHPS in pure culture [19]. Further, *B. wadsworthia*, a confirmed DHPS degrader in the human gut [15], is not widely distributed among mice and, if present, mostly derived from inoculation of a human strain or human feces transplants [19]. Among the DHPS degradation pathway-encoding taxa in the mouse microbiota, only *E. clostridioformis* was significantly enriched in the mouse gut content microcosms. While we showed that the murine *E. clostridioformis* strain YL32 ferments SQ to DHPS for growth, further degradation of DHPS to 3-HPS was not observed. So-called ’extended sulfoglycolytic pathways’ include genes for degradation of SQ and SQ degradation products such as DHPS or sulfolactate [83]. For example, *E. gilvus* contains the sulfofructose-transaldolase pathway and the bifurcated HpfGHD-HpfXYZ pathway for SQ and DHPS degradation [18]. *F. prausnitzii* is thought to also possess an extended sulfoglycolytic pathway, as its genome encodes the sulfofructose-transaldolase pathway along with a putative *hpfG* [17]. The HpfGHD pathway and the HpfXYZ pathway both do not produce H_2_S from DHPS [17,18]. The HpfGHD pathway does not conserve energy, but regenerates NAD^+^ [17], while ATP is generated in the bifurcated HpfXYZ pathway [18]. Our results indicate that DHPS is not a relevant growth substrate for the gut microbiota of conventional laboratory mice. Unlike the human gut, where SQ degradation proceeds beyond DHPS, SQ degradation stopped at the level of DHPS in the mouse gut and did not contribute to sulfide production. However, a previous study reported significant sulfide production after 168 h incubation of anoxic microcosms with feces of IL-10-deficient mice and 4 mM SQ, which indicated complete SQ degradation by the microbiota of this mouse model [82]. Microbiota composition was not analyzed in the fecal microcosms of IL-10-deficient mice. Yet, the discrepancy between the findings of this previous study and our study may be attributed to differences in microbial community structure and in incubation times. H_2_S-producing bacteria of the family *Desulfovibrionaceae* (genera *Taurinivorans*, *Desulfovibrio*, and *Mailhella*) seem to be generally more abundant in wild mice compared to laboratory mice [19,28,84]. Whether the absence of an active DHPS metabolism observed in the laboratory mice in our study reflects a host-specific microbial adaptation that is conserved across diverse wild and laboratory mouse lineages [85] or results from the specific diet and housing of laboratory mice remains unclear.

## Conclusions

Our work demonstrates that dietary SQ is exclusively degraded by the gut microbiota and not directly metabolized by the human and murine host. We observed the enrichment of commensal bacteria and significantly increased acetate and butyrate production in human fecal microcosms at high physiological SQ concentrations. These results align with key criteria for prebiotics [70]. SQ degradation by the gut microbiota of conventional wild type laboratory mice differed fundamentally from humans, primarily by the lack of DHPS degradation. Gnotobiotic animals colonized with defined consortia of microbial strains, including primary and secondary SQ degraders, may be better models to study effects of complete SQ degradation on the host [79,82]. Our study provides fundamental insight into SQ degradation in the mammalian gut, and has implications for potential application of SQDG-containing foods or SQ in dietary and prebiotic interventions.

## Declarations

### Ethics approval and consent to participate

Sampling and microbial analyses of feces from healthy human adults was approved by the ethics committee of the University of Vienna, Austria (reference numbers 00161 and 00714). Experiments with wild type mice were approved by the national authorities in Austria (Bundesministerium für Bildung, Wissenschaft und Forschung, 2022-0.745.974). Experiments with germ-free mice were approved by the local authorities in Germany (Regierung von Oberbayern, ROB-55.2-2532.Vet_02-19-53).

### Consent for publication

Not applicable.

### Availability of data and material

All sequence data generated in this project is available at NCBI under BioProject ID PRJNA1198716. The hidden Markov models developed in this project (Table S1) are available as Additional File 1.

## Funding

This research was funded by the Austrian Science Fund (FWF) [grant DOI 10.55776/DOC69 and 10.55776/COE7]. For open access purposes, the authors have applied for a CC BY public copyright license to any author-accepted manuscript version arising from this submission. BS received funding from the DFG (project no. 395357507 “SFB1371”), the European Research Council (EvoGutHealth, 865615), and the Germany Centre for Infection Research (project no. 503-5-7-06.712_00 and 503-5-7-06.709_00).

### Authors’ contributions

J.K. and A.L. conceived the study. J.K., B.T.H., A.S.W., S.B., M.L., and G.A. performed experiments and analyzed data. A.R., D.B., D.M., D.S., and B.S. contributed essential experimental infrastructure and analytical instrumentation, and expert advice. J.K., T.S.T., and B. H. performed bioinformatic analyses. J.K., B.T.H., A.S.W., S.B., T.S.T., M.M., and A.L. interpreted the data. J.K., M.M., and A.L. wrote the article. All authors revised and approved the manuscript.

## Acknowledgements

We thank the staff of the Joint Microbiome Facility for sequencing and Margarete Watzka for isotope ratio mass spectrometry measurements.

