## Supplementary material for "Sulfoquinovose is exclusively metabolized by the gut microbiota and degraded differently in mice and humans": Figure S

### Supplementary Figures

**Figure S1.** Sulfur metabolites and short-chain fatty acid dynamics in 24 h human fecal microcosm incubations with different SQ concentrations.

**Figure S2.** Enriched ASVs in SQ-amended human fecal microcosm incubations.

**Figure S3.** Isethionate production in mouse gut content microcosms amended with 10 mM SQ (Experiment 1).

**Figure S4.** Changes in total bacterial and archaeal abundance in mouse gut content microcosms amended with 10 mM SQ or 10 mM DHPS.

**Figure S5.** Enriched ASVs in mouse gut content microcosm incubations amended with 10 mM SQ (Experiment 1).

**Figure S6.** Enriched ASVs in mouse gut content microcosm incubations amended with 10 mM SQ (Experiment 2).

**Figure S7.** Fecal SQ and DHPS concentrations in conventional mice after oral SQ supplementation as measured by capillary electrophoresis.

**Figure S8.** *Faecalibaculum rodentium* did not degrade SQ in both rich and diluted rich medium.

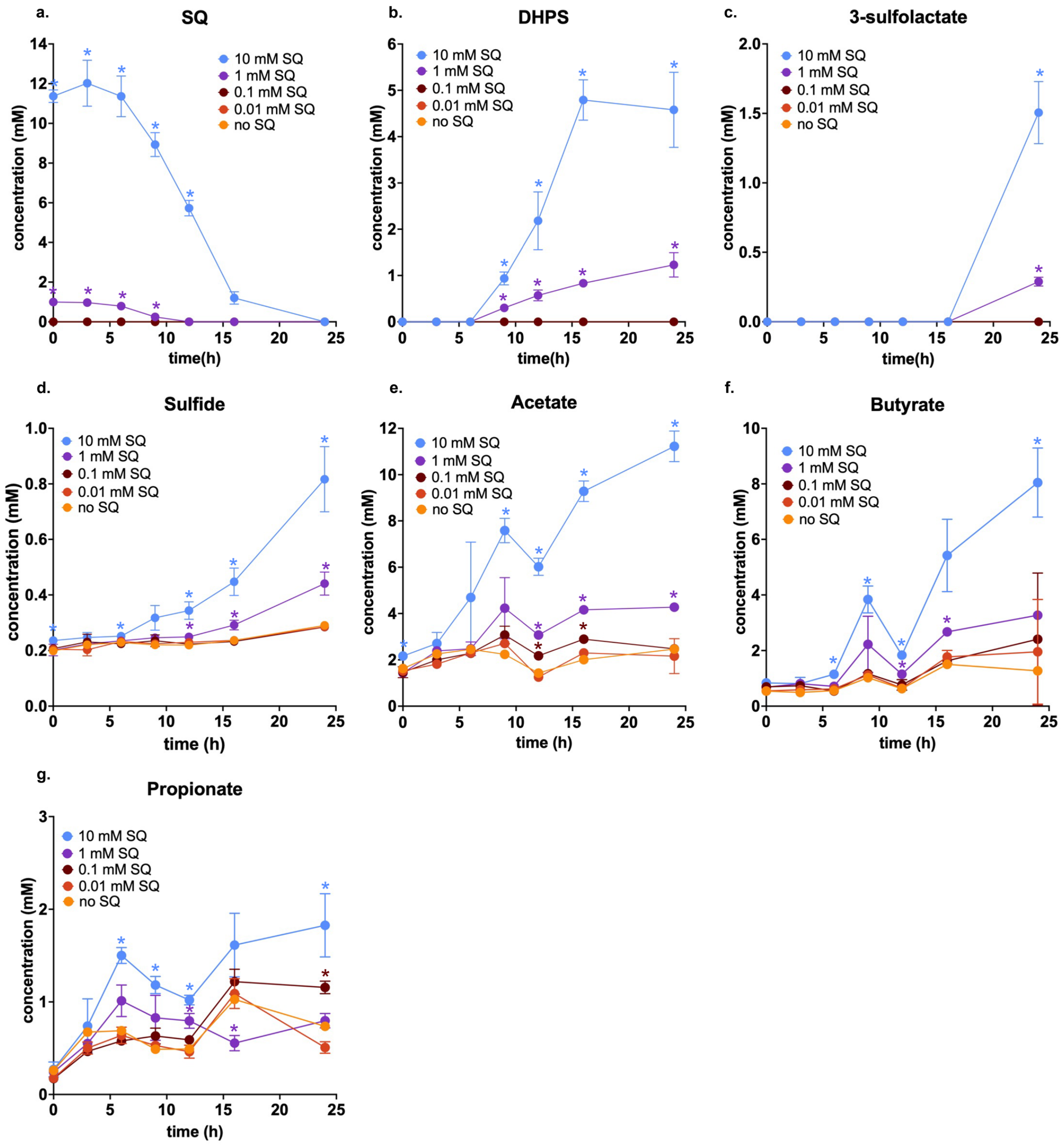


**Figure S1. Sulfur metabolites and short-chain fatty acid dynamics in 24 h human fecal microcosm incubations with different SQ concentrations.** Dynamics of the sulfur metabolites SQ (**a**), DHPS (**b**), 3-sulfolactate (**c**), sulfide (**d**), and the short-chain fatty acids acetate (**e**), butyrate (**f**), and propionate (**g**) in incubations with different concentrations of SQ (10 mM, 1 mM, 0.1 mM, 0.01 mM), as determined by capillary electrophoresis. Data points represent the average of triplicate incubations and the error bars represent one standard deviation. Only one unamended control incubation was performed. Coloured asterisks indicate significant differences between treatment groups and the pooled control group (unamended control + 0.01 mM SQ treatment), at individual time points, based on ANOVA with Dunnett's post hoc test. Adjusted *p*-values below 0.05 are considered significant. SQ, sulfoquinovose; DHPS, 2,3-dihydroxypropane-1-sulfonate.

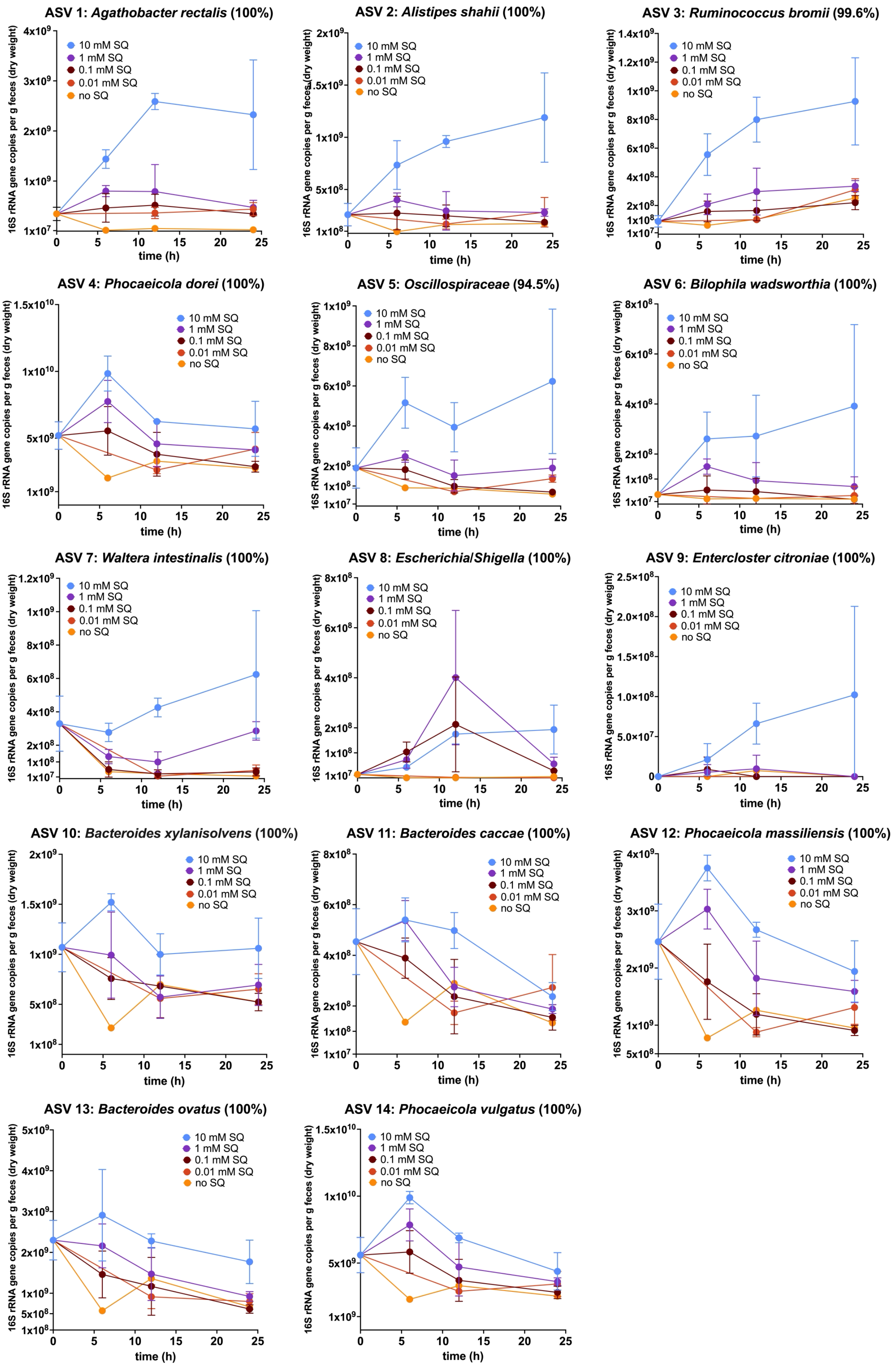
ASV 1: Agathobacter rectalis (100%)
ASV 2: Alistipes shahii (100%)
ASV 3: Ruminococcus bromii (99.6%)
ASV 4: Phocaeicola dorei (100%)
ASV 5: Oscillospiraceae (94.5%)
ASV 6: Bilophila wadsworthia (100%)
ASV 7: Waltera intestinalis (100%)
ASV 8: Escherichia/Shigella (100%)
ASV 9: Enterocloster citroniae (100%)
ASV 10: Bacteroides xylanisolvens (100%)
ASV 11: Bacteroides caccae (100%)
ASV 12: Phocaeicola massiliensis (100%)
ASV 13: Bacteroides ovatus (100%)
ASV 14: Phocaeicola vulgatus (100%)
10 mM SQ
1 mM SQ
0.1 mM SQ
0.01 mM SQ
no SQ
16S rRNA gene copies per g feces (dry weight)
time (h)

**Figure S2. Enriched ASVs in SQ-amended human fecal microcosm incubations.** Changes in absolute abundance of the fourteen significantly enriched ASVs in incubations amended with 10 mM SQ, 1 mM SQ and/or 0.1 mM SQ, as identified via differential abundance analysis. ASVs are shown in order of their average increase in absolute abundance after 24 h of incubation. Points represent averages of three replicates, except for the unamended control, of which only one replicate was sequenced. Error bars represent one standard deviation. Taxonomic classification and the 16S rRNA gene sequence similarity to the closest relative (in brackets) is shown for each ASV. ASV, amplicon sequencing variant; SQ, sulfoquinovose.

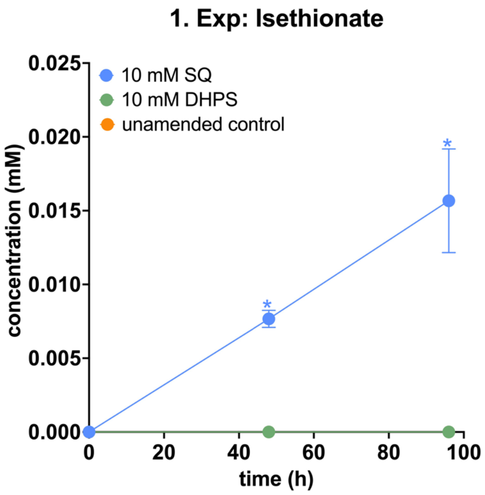


**Figure S3. Isethionate production in mouse gut content microcosms amended with 10 mM SQ (Experiment 1).** Small amounts of isethionate were only produced in SQ-amended microcosms, as determined via liquid chromatography-mass spectrometry. Lines show the average of triplicate measurements and error bars represent one standard deviation. Asterisks show significant differences between treatment groups and the unamended control at individual time points determined using ANOVA with Dunnett's post hoc test. Adjusted *p*-values below 0.05 were considered significant. Exp, experiment; SQ, sulfoquinovose; DHPS, 2,3-dihydroxypropane-1-sulfonate.

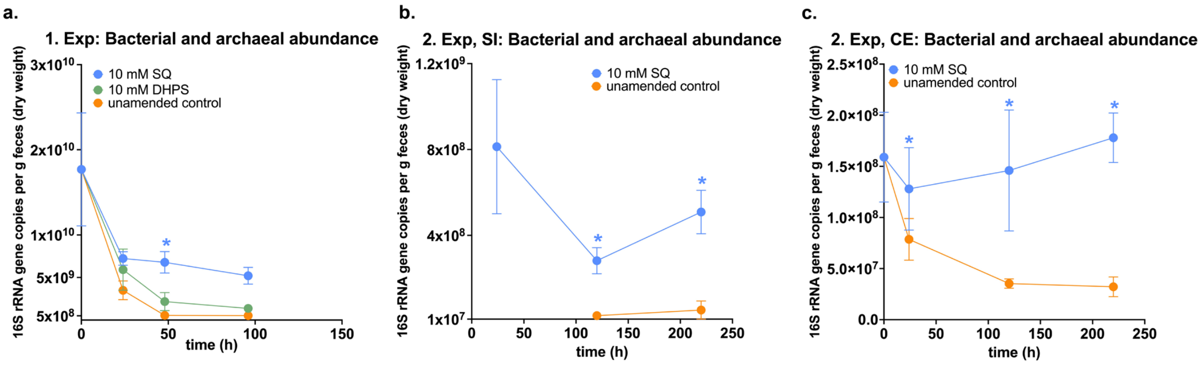


**Figure S4. Changes in total bacterial and archaeal abundance in mouse gut content microcosms amended with 10 mM SQ or 10 mM DHPS.** Dynamics of bacterial and archaeal 16S rRNA gene copy numbers in SQ-amended mouse gut content microcosms from pooled gut contents (**a**), small intestinal contents (**b**), and cecal contents (**c**). In **b**., quantification of samples from time point zero h failed. All lines show the average of triplicate measurements and error bars represent one standard deviation. In **a**., coloured asterisks show significant differences between treatment groups and their respective control groups without SQ at individual time points, as determined using ANOVA with Dunnett's post hoc test. In **b**. and **c**., significant differences between SQ-amended incubations and the unamended control were determined at each time point via t-test and are depicted as asterisks. Adjusted *p*-values below 0.05 were considered significant. Exp, experiment; SQ, sulfoquinovose; DHPS, 2,3-dihydroxypropane-1-sulfonate; SI, small intestine; CE, cecum.

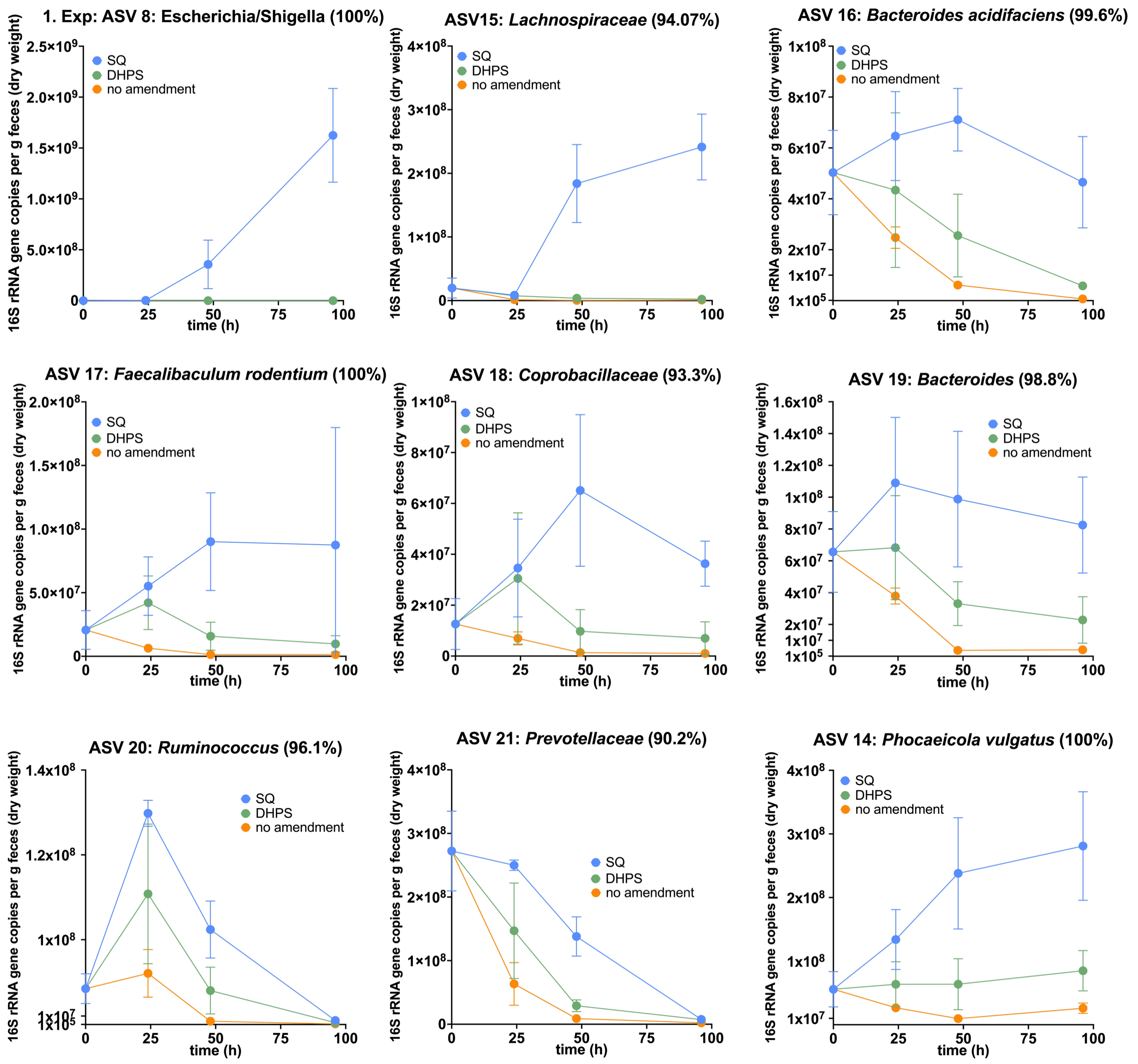


**Figure S5. Enriched ASVs in mouse gut content microcosm incubations amended with 10 mM SQ (Experiment 1).** Changes in absolute abundance of the nine significantly enriched ASVs in incubations amended with 10 mM SQ, as identified via differential abundance analysis. No ASVs were significantly enriched in DHPS-amended incubations. ASVs are shown in order of their $\log_2$ fold change between SQ-amended incubations and the unamended control. Data points represent averages of three replicates and error bars represent one standard deviation. Taxonomic classification and the 16S rRNA gene sequence similarity to the closest relative (in brackets) is shown for each ASV. ASV, amplicon sequencing variant; SQ, sulfoquinovose; DHPS, 2,3-dihydroxypropane-1-sulfonate.

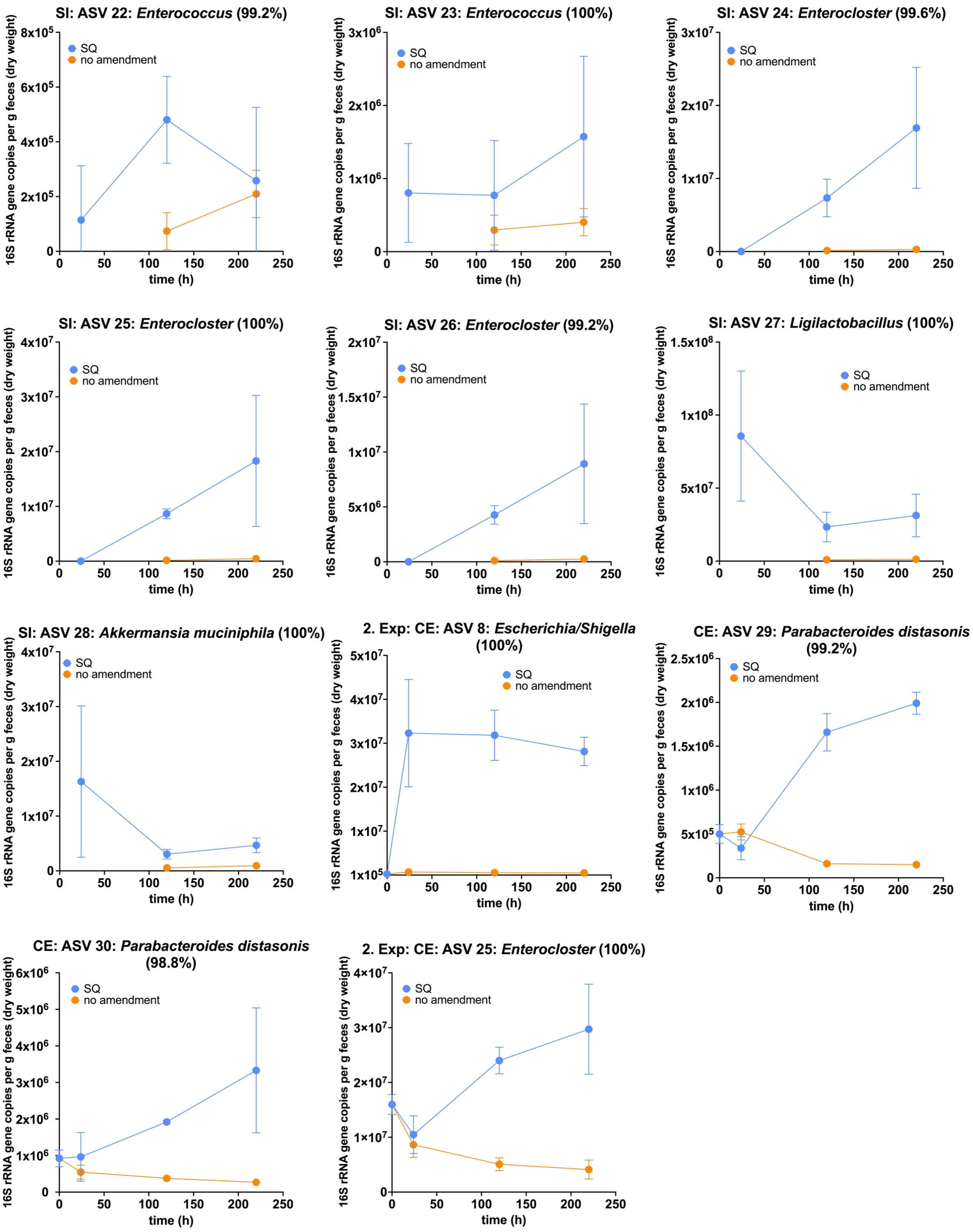


**Figure S6. Enriched ASVs in mouse gut content microcosm incubations amended with 10 mM SQ (Experiment 2).** Changes in absolute abundance of significantly enriched ASVs in SQ-amended incubations, with seven enriched ASVs in incubations with small intestinal contents and four enriched ASVs in incubations with cecal contents, as identified via differential abundance analysis. *Enterocloster* ASV 25 was enriched in the SQ-amended small intestinal content incubations and cecal content incubations. ASVs are shown in order of their $\log_2$ fold change between SQ-amended incubations and the unamended control. Points represent averages of three replicates. Error bars represent one standard deviation. Taxonomic classification and the 16S rRNA gene sequence similarity to the closest relative (in brackets) is shown for each ASV. SI, small intestine; CE, cecum; ASV, amplicon sequencing variant; SQ, sulfoquinovose.

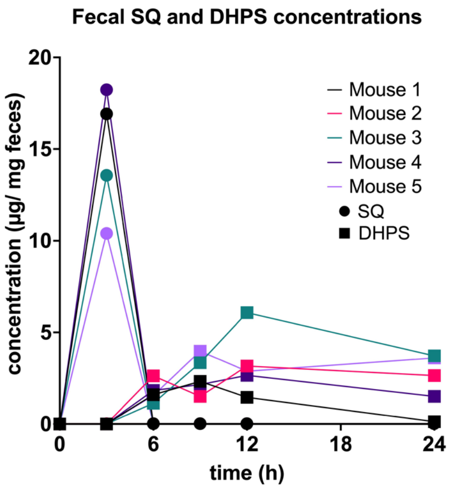


**Figure S7. Fecal SQ and DHPS concentrations in conventional mice after oral SQ supplementation as measured by capillary electrophoresis.** Ten mg SQ was metabolized within six hours to DHPS in conventional C57BL/6 mice. DHPS was detected between 6 h and 24 h post-gavage. Metabolite dynamics were additionally measured using liquid chromatography-mass spectrometry (Figure 5 b, Table S4). The discrepancies between the two methods are primarily due to (i) the lower sensitivity and specificity of capillary electrophoresis, as well as technical variability between the methods, and (ii) the fact that only a subset of the samples analyzed by capillary electrophoresis were also examined using liquid chromatography–mass spectrometry. SQ, sulfoquinovose; DHPS, 2,3-dihydroxypropane-1-sulfonate.

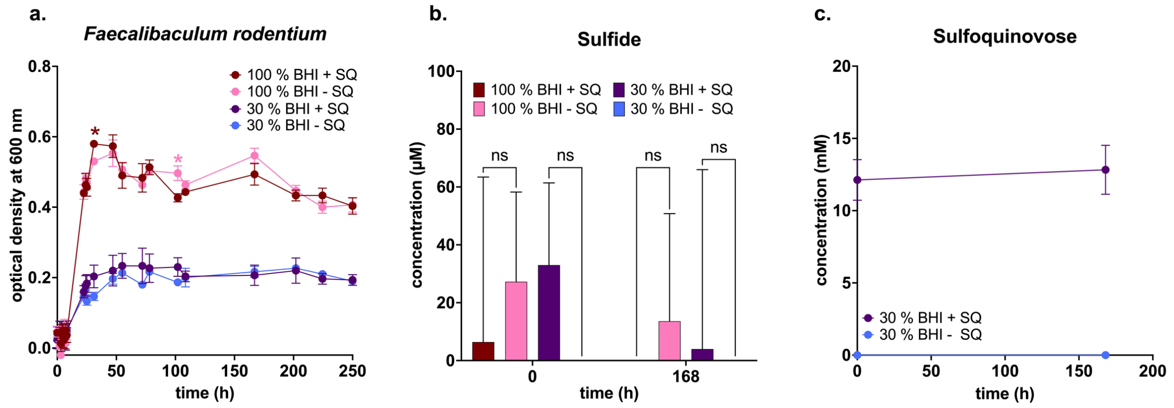


**Figure S8.** ***Faecalibaculum rodentium*** **did not degrade SQ in rich medium and diluted rich medium.** Growth dynamics of *F. rodentium* in undiluted (100%) and diluted (30%) BHI medium with and without 10 mM SQ (**a**). Sulfide (**b**) and sulfoquinovose (**c**) concentrations in *F. rodentium* cultures with or without SQ. SQ concentrations were measured in diluted BHI medium via capillary electrophoresis. SQ addition did not lead to growth stimulation, sulfide production or SQ degradation. Data points in **a**. and **c**. show averages of triplicate cultures with error bars representing one standard deviation. In **a**., coloured asterisks show significant differences between treatment groups and their respective control groups without SQ at individual time points. Bars in **b**. represent average sulfide concentrations and error bars represent one standard deviation. Values that were negative after subtraction of the blank were set to zero. Brackets show results of pairwise comparisons between treatment groups and their respective unamended control. Significant differences were determined using ANOVA with Tukey's post hoc test. In **c**., SQ concentrations at 0 h were compared to SQ concentrations at 168 h via t-test. Adjusted *p*-values below 0.05 were considered significant. SQ, sulfoquinovose; BHI, brain-heart infusion; ns, not significant.
